# The knockout of the HMG domain of the porcine SRY gene causes sex reversal in gene-edited pigs

**DOI:** 10.1101/617589

**Authors:** Stefanie Kurtz, Andrea Lucas-Hahn, Brigitte Schlegelberger, Gudrun Göhring, Heiner Niemann, Thomas C. Mettenleiter, Björn Petersen

## Abstract

The sex-determining region on the Y chromosome (SRY) is thought to be the central genetic element of male sex development. Mutations within the SRY gene are associated with a male-to-female sex reversal syndrome in humans and other mammalian species such as mice and rabbits. However, the underlying mechanisms are largely unknown. To understand the biological function of the SRY gene, a site-directed mutational analysis is required to investigate associated phenotypic changes at the molecular, cellular and morphological level. In our study, we successfully generated a knockout of the porcine SRY gene by microinjection of two clustered regularly interspaced short palindromic repeats (CRISPR) – associated protein - 9 nuclease (Cas9) ribonucleoprotein (RNP) complexes targeting the centrally located “high mobility group” (HMG) domain of the SRY gene. Mutations within this region resulted in the development of complete external and internal female genitalia in genetically male pigs. The internal female genitalia including uteri, ovaries, and oviducts, revealed substantial size differences in 9-months old SRY-knockout pigs compared to age-matched female wild type controls. In contrast, a deletion within the 5’ flanking region of the HMG domain was not associated with sex reversal. Results of this study demonstrates for the first time the central role of the HMG domain of the SRY gene in male sex determination in pigs. Moreover, quantitative analysis by digital PCR revealed evidence for a duplication of the porcine SRY locus. Our results pave the way towards the generation of boars exclusively producing phenotypically female offspring to avoid surgical castration without anesthesia in piglets. Moreover, the study establishes a large animal model that is much more similar to humans in regard of physiology and anatomy and pivotal for longitudinal studies.

## 2 Introduction

In mammals, the male and female sex are determined by the presence or absence of the Y chromosome (1). The sex-determining region on the Y chromosome (SRY) is located on the short arm of the Y chromosome and is presumed to be critical for sex determination during embryogenesis (2, 3). In pigs, the SRY gene consists of a single exon, with an open reading frame of 624 bp representing 206 amino acids and encodes for the testis-determining transcription factor (TDF). It is expressed in the male genital ridge at the time of sex determination (4). The porcine SRY gene is first expressed on day 21 post coitum (p.c.) with highest expression levels between day 21 and 23 p.c. Shortly after onset of SRY expression, testis formation can be histologically determined between day 24 to 27 p.c. (5, 6). Accordingly, the SRY gene is assumed to serve as the master regulator causing the formation of primary precursor cells of tubuli seminiferi leading to the development of testicles from undifferentiated gonads (7). However, it is still unknown whether the SRY gene is the only sex-determining gene on the Y chromosome or if other genes such as SOX9 (8-10) and SOX3 (11) are involved as well.

In previous studies in mice (12) and rabbits (13), the SRY gene was knocked out using different target regions. Both, the knockout of 92 % of the murine SRY gene by TALENs, or the CRISPR/Cas-mediated knockout of the Sp1-DNA-binding sites of the rabbit SRY gene caused sex reversal. Nevertheless, sequence divergence of the SRY gene between mammalian species limited its direct structural and functional comparison and the investigation of mammalian sex determination. So far, analysis of the SRY gene has almost exclusively been done in small animals, mostly mice, and knowledge about the SRY gene in large animal species, especially the porcine SRY gene is scarce.

The goal of the present study was to characterize the porcine SRY gene and its HMG domain in male sex determination by knocking out different target sites of the porcine SRY gene via intracytoplasmic microinjection of two CRISPR/Cas9 RNPs or cell transfection followed by somatic cell nuclear transfer (SCNT) (Fig. 1). The generation of SRY-knockout pigs give insights into the biological function of the SRY gene in a large animal species. While, the murine SRY gene shows only 75% similarity to the human SRY gene, the porcine and human SRY genes are closely related (∼85 % amino acid homology) and show similar expression profiles (5, 14). Therefore, a knockout of the highly conserved HMG domain in the porcine model may pave the way for a suitable large animal model for the human male-to-female sex reversal syndrome.

**Figure 1.**
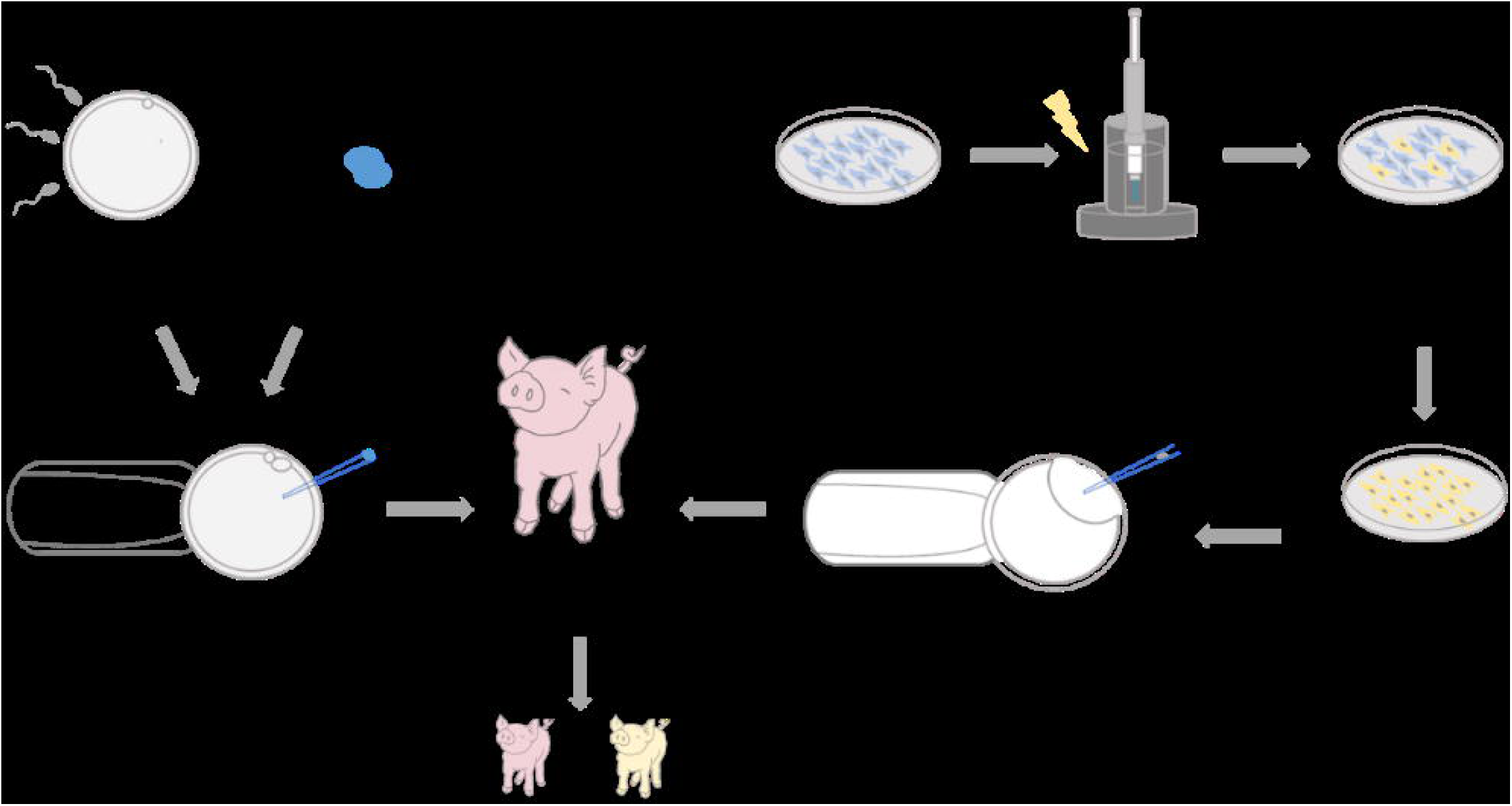
Schematic illustration of the experimental design to generate SRY-KO pigs (XY^SRY-^) by either intracytoplasmic microinjection of two CRISPR/Cas9 RNP complexes into IVF-produced zygotes or somatic cell nuclear transfer (SCNT). Embryos were surgically transferred into hormonally synchronized recipients, and the offspring were analyzed pheno- and genotypically.

## 4 Results

### Production of SRY-knockout pigs

To generate SRY-knockout (SRY-KO) pigs, a deletion of approx. 300bp, encompassing the HMG domain in the porcine SRY gene was introduced (Fig. 2a). Thirty-one and thirty-two embryos derived from intracytoplasmic microinjection of gRNA SRY_1 and SRY_3 into IVF-produced zygotes were surgically transferred into three synchronized sows. Two recipients went to term and delivered twelve healthy piglets with female phenotypes (Tab. 1/Fig. 3). Three of the piglets showed a deletion of approx. 300 bp within the HMG domain of the SRY gene (Fig 4). Sequencing of the target region revealed frameshift mutations of – 266 bp in piglet 715/2, and – 292 bp in piglet 715/7. Two different genetic modifications, including a deletion of 298 bp and an indel formation with a deletion of 298 bp and an insertion of 1 bp were detected in piglet 714/1 (Fig. 5).34 Furthermore, analysis of six Y chromosome specific genes (KDM6A, TXLINGY, DDX3Y, CUL4BY, UBA1Y and UTY) demonstrated a male genotype and successful sex reversal in these piglets (Fig. S1, Tab. S1). To ultimately confirm the male genotype of these piglets (715/2, 715/7 and 714/1), cells from ear tissue were karyotyped detecting the Y chromosome in all three piglets (Fig. 6, Fig. S2). No chromosomal abnormalities were observed in the sex-reversed pigs 715/2 and 715/7, while piglet 714/1 revealed an inversion of chromosome 7 (Fig. S2). The origin of this clonal cytogenetic aberration remains unclear. It may not necessarily be related to the CRISRP/Cas system because no off-target event was found on chromosome 7. In total, 34 potential off-target sites within the porcine genome were identified (http://crispor.tefor.net/). We designed primers for the top ten off-target sites for each gRNA (Tab. S2/S3). In one off-target site for gRNA_SRY1 and three off-target sites for gRNA_SRY3 PCR amplification following Sanger sequencing was not possible. Overall, no off-target events were observed (Fig. S3-S5). All SRY-KO pigs developed normally without any health impairment (Fig. S6, Tab. S4).

**Figure 2.**
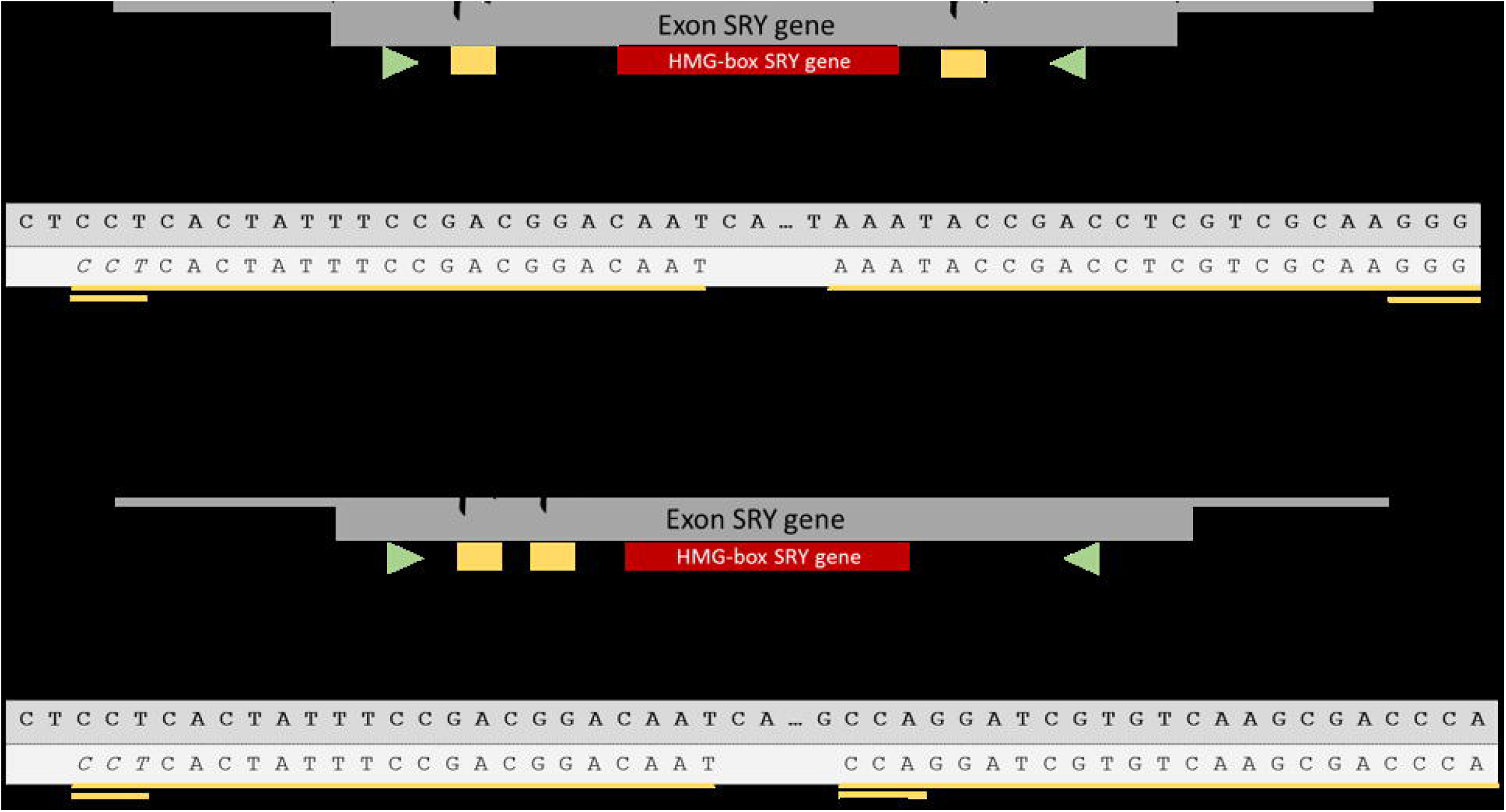
**a** Location of two sgRNA target sites (yellow underlined) flanking the HMG-box (red box) of the SRY gene. **b** Schematic illustration showing the guide RNAs (yellow underlined) targeting an approx. 72 bp segment in the 5’ flanking region of the HMG domain (red box) of the SRY gene. Primer amplifying the SRY exon are indicated with green arrows.

**Table 1.** Results of the embryo transfer of microinjected zygotes into recipients. Three of twelve piglets showed a sex reversal with a female phenotype and a male genotype.

|  |  |  |  |  |  |
| --- | --- | --- | --- | --- | --- |
| 8018 | 32 | - | - | - | - |
| 715 | 31 | + | 11 | 2 | 2 |

**Figure 3.**
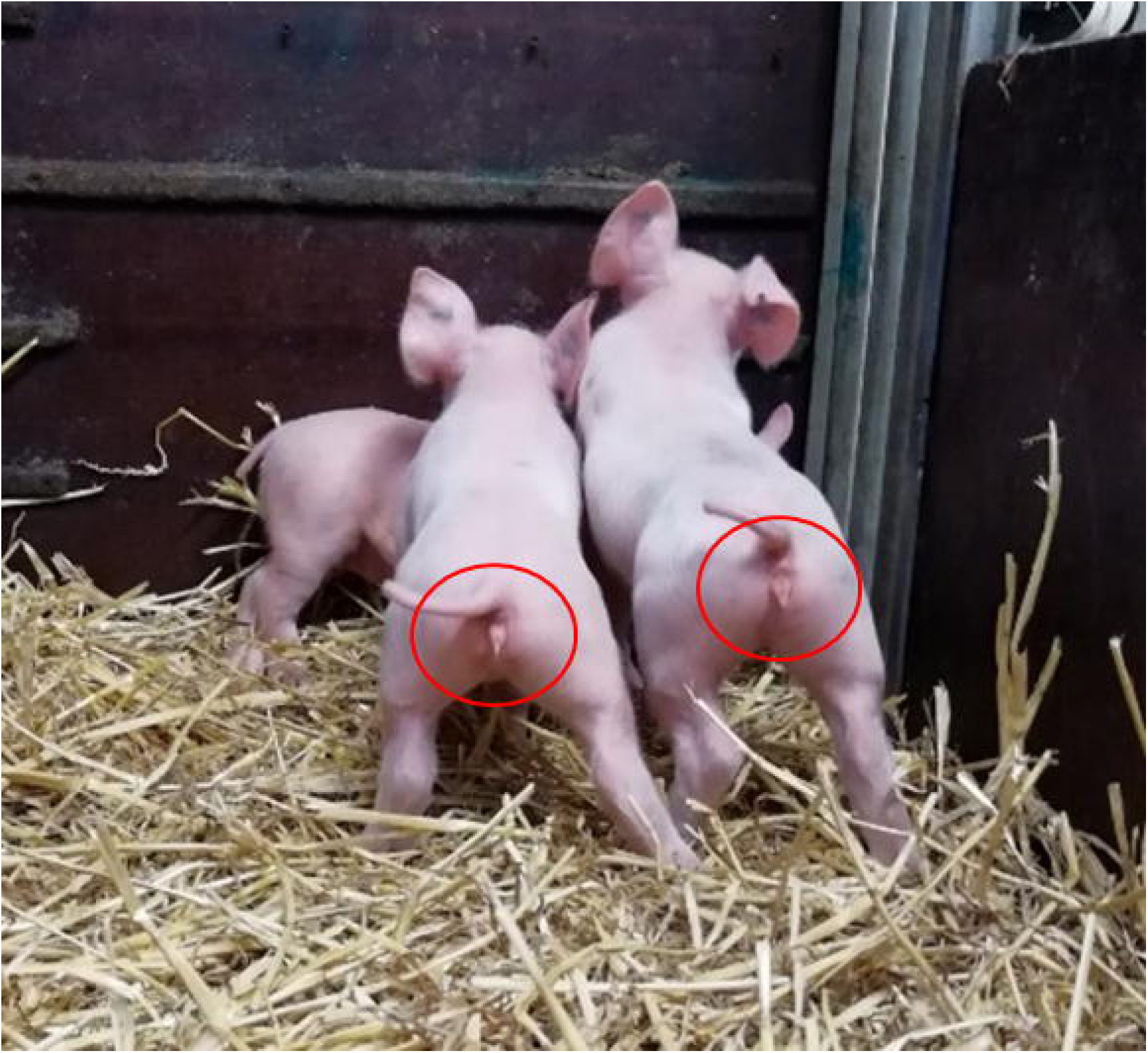
Twelve healthy piglets were born after cytoplasmic microinjection of two CRISPR/Cas9 RNP complexes into IVF-produced zygotes and surgical embryo transfer. Three of the piglets showed complete female external genitalia. The deletion of the SRY gene had no effect on growth rate compared to wild type. All piglets developed normally.

**Figure 4.**
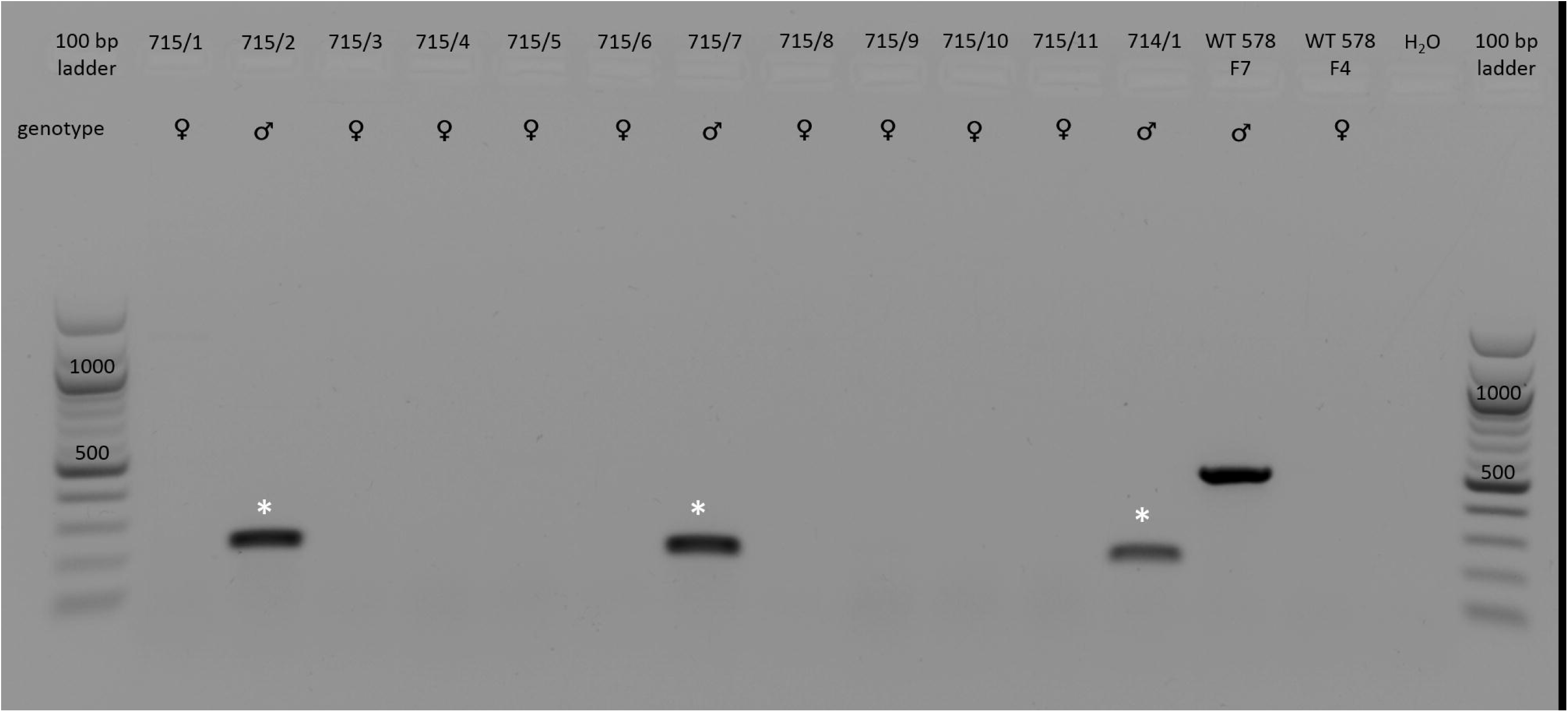
PCR-based detection of the mutated SRY gene in piglets (714/1 and 715/1-11) generated via microinjection of CRISPR/Cas9 RNP complexes. Three piglets (715/2, 715/7 and 714/1, indicated with white asterisk) showed deletions of approx. 300 bp within the SRY gene compared to a male wild type control (WT 578 F7). The male WT control showed an expected band of ∼500 bp. The female WT control (WT 578 F4) is negative, as expected for the SRY gene.

**Figure 5.**
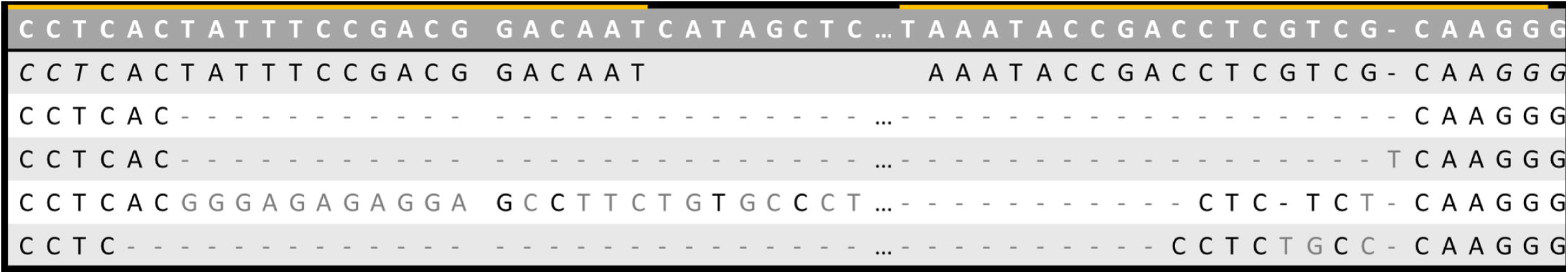
Sanger sequencing of the purified PCR product of the SRY-KO piglets (715/2, 715/7 and 714/1) showed genetic modifications within the SRY locus. Piglet 715/7 displayed a deletion of 292bp and piglet 715/2 of 266bp. Piglet 714/1 showed two different mutations with a deletion of 298bp and an indel formation of −298bp and +1bp.

**Figure 6.**
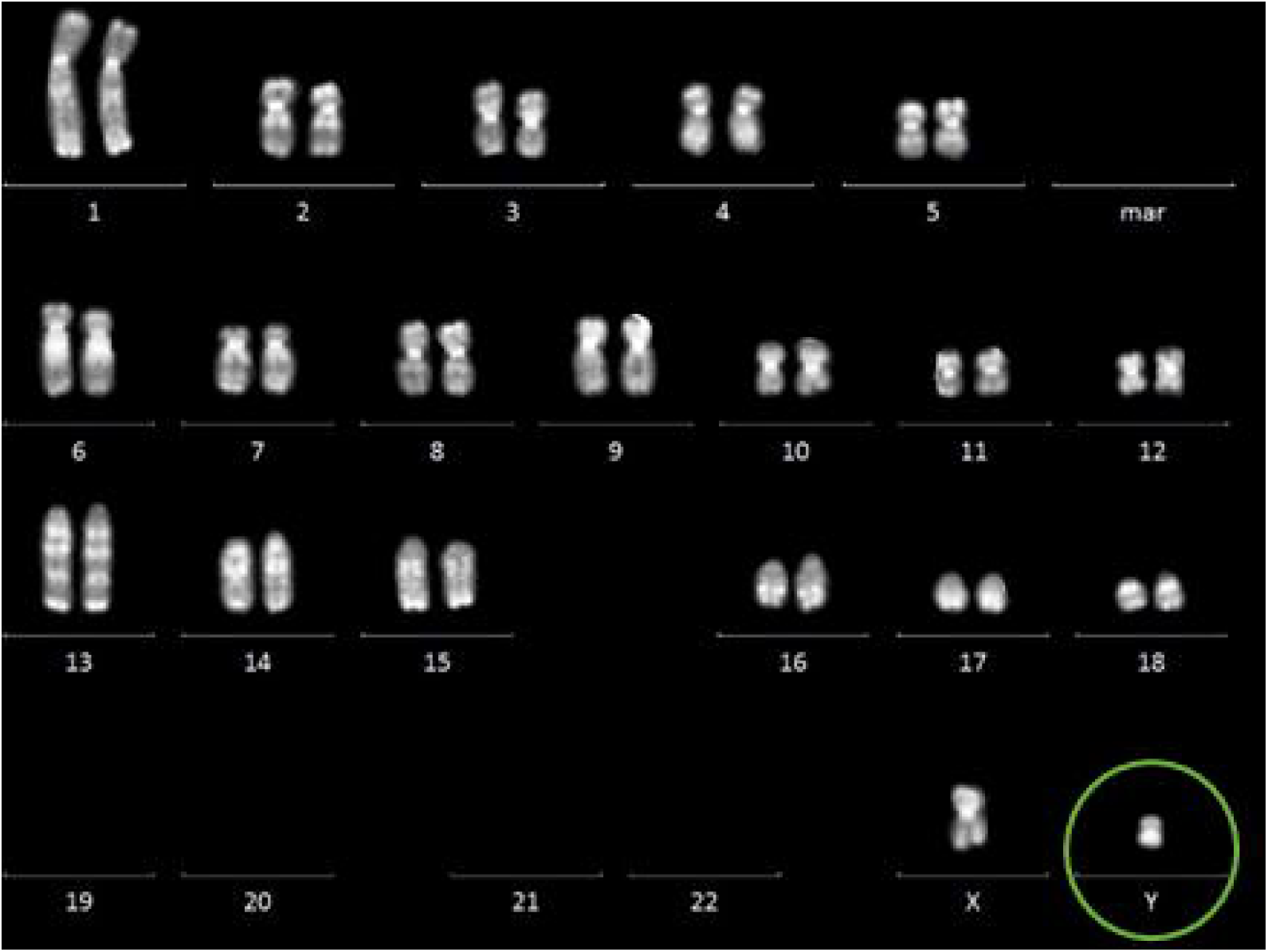
Karyotyping of cells from the SRY-KO piglet 715/2 confirmed the male genotype of this piglet. The karyotypes of piglet 715/7 and 714/1 are shown in the Supplements Fig. 6.

In a second approach, one piglet generated via intracytoplasmic microinjection of gRNAs SRY_1 and SRY_2 targeting the 5’ flanking region of the HMG domain (Fig. 2b) was born with an in-frame mutation of – 72 bp on the SRY locus, which did not lead to sex reversal (Fig. S7).

### External and internal genitalia of the SRY-KO pigs

We investigated the external and internal genitalia of the SRY-KO pigs. Age-matched wild type (WT) females from conventional artificial insemination and female littermates of the SRY-KO pigs produced by microinjection served as controls. At the age of 34 days, the external genitalia of the SRY-KO piglets were similar to the external genitalia of female littermates and WT controls. To further investigate the internal genitalia, the ovaries, oviducts and uteri of the 34 days old SRY-KO piglets and female controls were prepared. The SRY-KO piglets had complete female internal genitalia, including ovaries, oviducts and uteri that were similar to that of age matched WT females (Fig. S8). Moreover, histological analysis of the inner structure of the ovaries revealed no alteration in these young piglets (Fig. S9).

However, substantial size differences of the female genitalia were obvious in 9-months old SRY-KO pigs compared to age-matched wild type controls (Fig. 7), with gene-edited animals showing a substantially smaller genital tract. The SRY-KO pigs were not observed in heat, even after three consecutive treatments of 1,000 IU PMSG (Pregmagon^®^, IDT Biologika) followed 72 hours later by an intramuscular injection of 500 to 1,000 IU hCG (Ovogest^®^300, MSD Germany) to induce estrus. Histological analysis of the ovaries of 9-months old SRY-KO pigs revealed a high amount of loose connective tissue (Fig. S10). These results provided evidence that the SRY-KO caused sex reversal in genetically male pigs with the development of female external and internal genitalia, which lends further support to the central role of the SRY gene in male sex determination during porcine embryogenesis. Re-cloning of piglet 715/2 led to seven sex-reversed piglets and demonstrated unequivocally that the elaborated strategy described in this study can be used to successfully produce sex-reversed pigs (Fig. S11 to S14).

**Figure 7.**
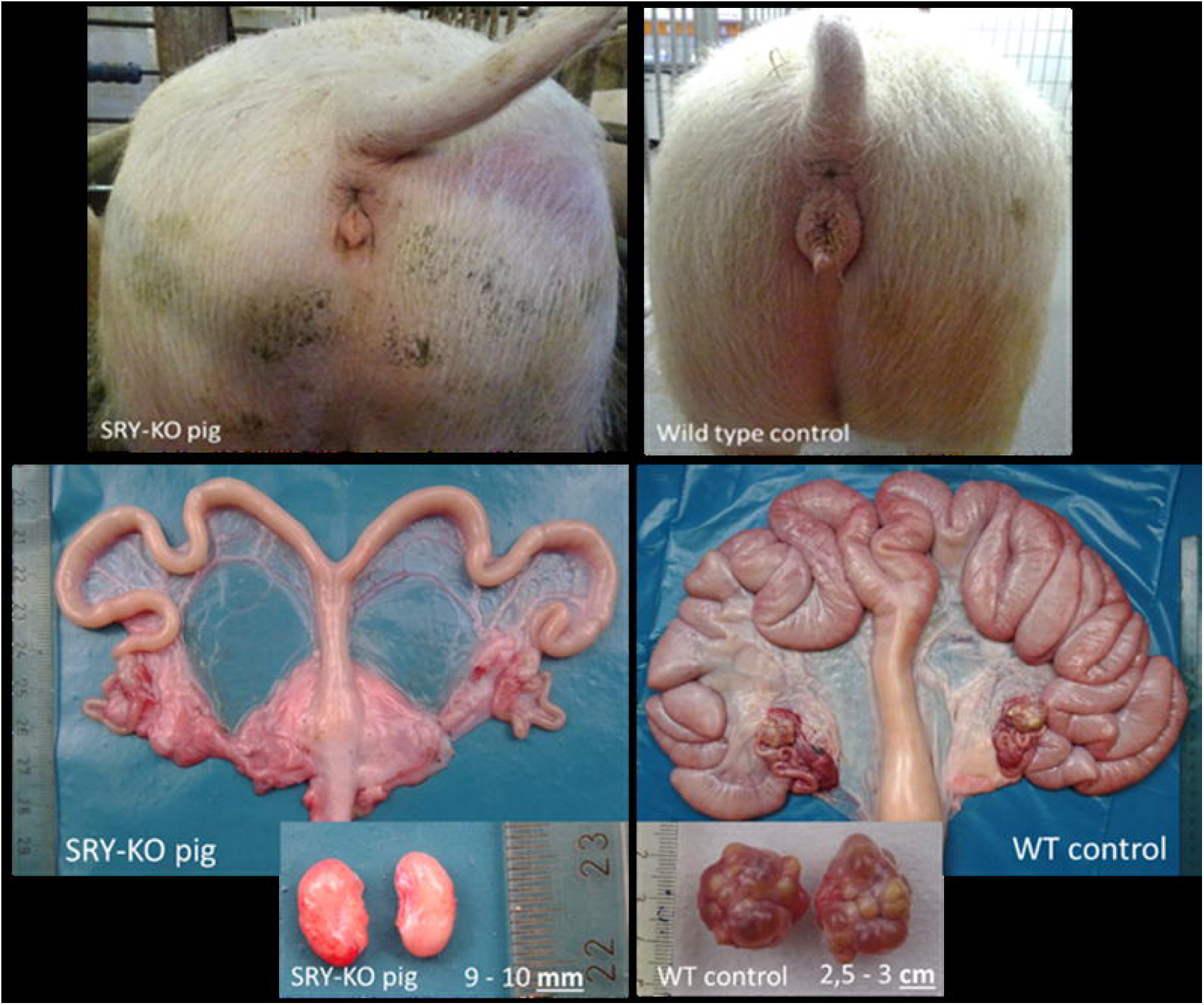
The uteri and ovaries of the 9-months old SRY-KO, XY pig (714/1) and the age-matched WT,XX piglet (control from same litter). a Substantial size differences were displayed in the 9-months old SRY-KO pig compared to the female wild type control. b The ovaries in the 9-month old SRY-KO, XY pig were significant smaller than the ovaries of the WT, XX pig and showed no follicles.

### Duplication of the porcine SRY gene

Investigation of the HMG domain of the porcine SRY gene revealed one pig derived from intracytoplasmic microinjection (714/1) that displayed two different genetic modifications within the SRY locus (Fig. 5). Whether these modifications originated from mosaicism caused by the microinjection or a duplication of the SRY locus was further analyzed. Analysis of different organ samples (liver, heart, colon, kidney, spleen and lung) of piglet 714/1 revealed the same two genetic modifications in all samples, arguing against mosaicism but for consideration of a duplicated SRY locus. Whole genome sequencing using the USB-connected, portable Nanopore sequencer MinION (Oxford Nanopore Technologies) and digitalPCR (QuantStudio^®^3D, ThermoFisher Scientific) was performed to check for a possible SRY gene duplication. Due to the high number of repetitive genes on the Y chromosome, Nanopore technology was used to sequence large DNA fragments. However, only one contig similar to the SRY sequence could be found with the assembled reads of the Nanopore Sequencing data. Even in a broad range, the sequence in the flanking regions of the SRY locus was nearly identical hampering the analysis of the duplicated SRY gene when using only the assembled Nanopore Sequencing data.

To further verify the duplication of the SRY gene, genomic DNA was analyzed by digital PCR. Three targets, including the monoallelic SRY and KDM6A genes on the Y chromosome, and the biallelic GGTA (galactosyltransferase) gene on chromosome 1 were selected for direct comparison of their copy numbers. The copy numbers of GGTA1 were set to two (biallelic), whereas the KDM6A and SRY genes were quantified in relation to the GGTA1 gene. In a first trial, comparison of the copy numbers of the KDM6A and GGTA1 genes revealed a 2-fold lower copy number of the monoallelic KDM6A compared to the biallelic GGTA1 in a male wild type control (WT 7214 F2), as expected. In contrast, the normally monoallelic SRY gene exhibited a similar calculated copy number as the biallelic GGTA1 gene (Fig. 8), indicating a duplication of the SRY gene.

**Figure 8.**
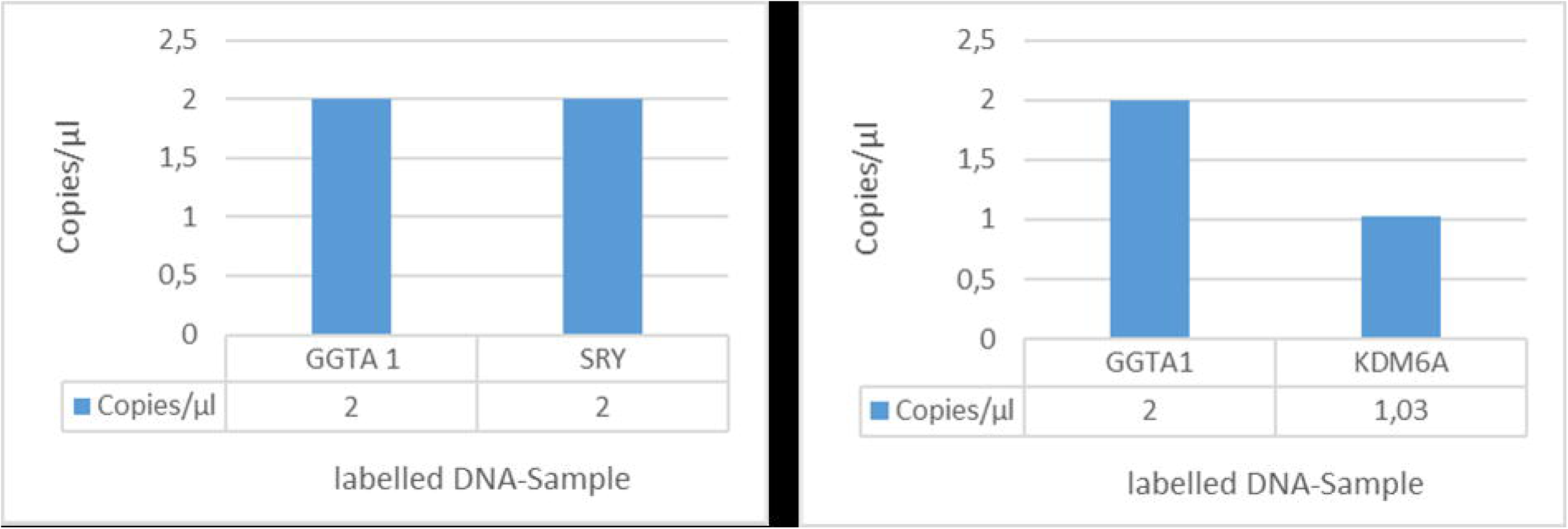
Schematic diagram of the first trail of dPCR. The dPCR biplex assay revealed a two fold lower copy number of the monoallelic KDM6A gene compared to the biallelic GGTA1 gene, as expected. A similar copy number of the monoallelic SRY gene compared to the biallelic GGTA1 gene indicated a duplication of the SRY locus.

A second approach confirmed these findings by comparison of the copy numbers of the SRY and GGTA1 gene in pigs with a complete SRY-KO (714/1, 715/2, 715/7), an incomplete SRY-KO (713/1) and wild type controls (Fig. 9). As expected, no signal for SRY was detected in pigs with a complete SRY-KO. For the incomplete SRY-KO, a piglet derived from intracytoplasmic microinjection of plasmids SRY_1 and SRY_3 was used that showed two genetic modifications, i.e. a 3 bp and a 298 bp deletion, within the SRY locus (Fig. S15/S16). Analysis of several organ samples (liver, heart, colon, spleen, kidney, epididymis, testis and lung) revealed the same genetic modification in all organs, indicating that mosaicisms was highly unlikely (Fig. S17). In this piglet, dPCR showed a 50 % reduced copy number of the SRY gene compared to the GGTA1 gene. The SRY probe bound to the SRY locus with the smaller deletion of 3 bp that did not interfere with the SRY assay and thereby indicated a duplication of the SRY locus. As mentioned above, a similar copy number of the monoallelic SRY gene compared to the biallelic GGTA1 gene was detected in wild type controls (Fig. 9).

**Figure 9.**
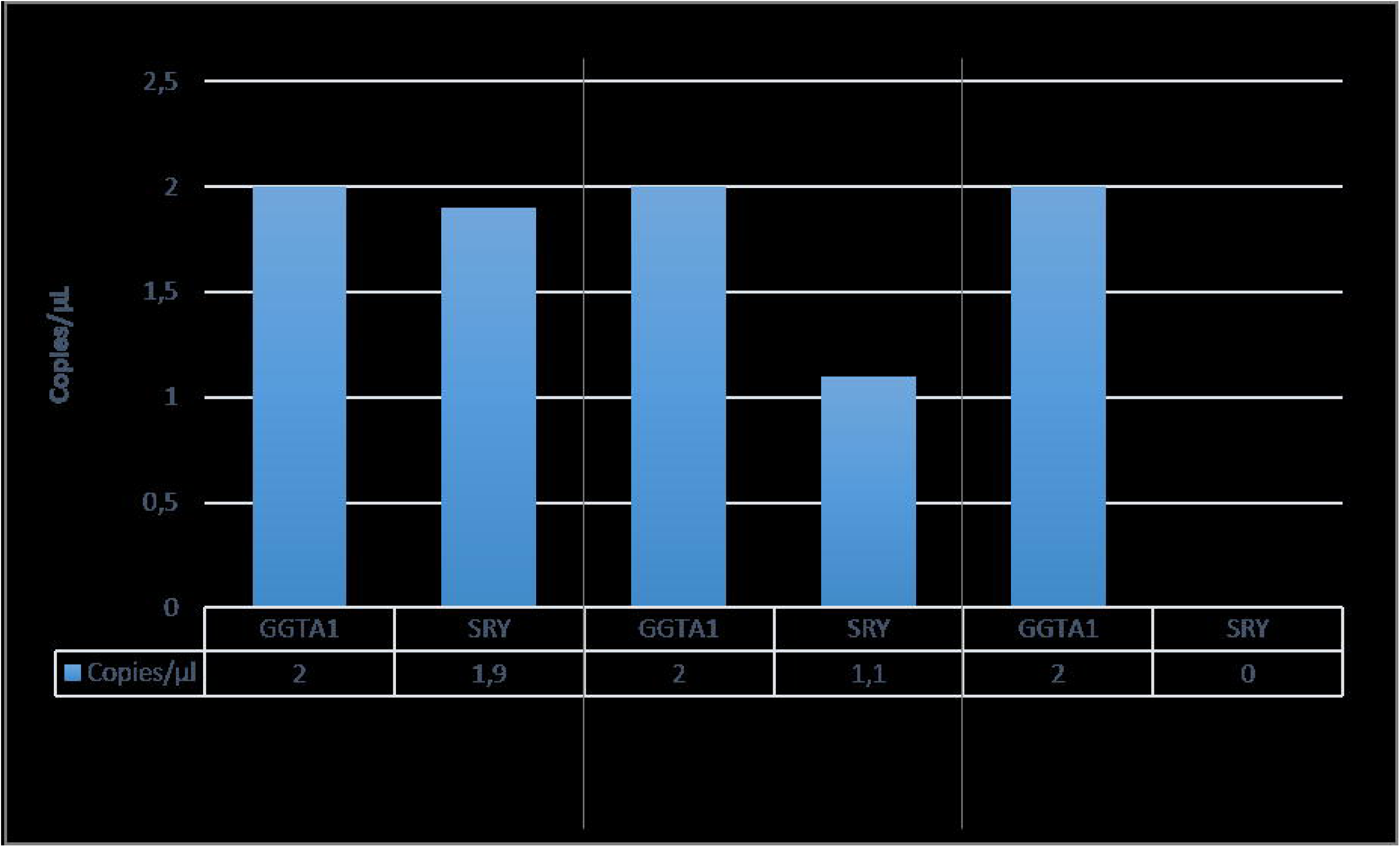
Schematic diagram of the second trail of dPCR. The dPCR biplex assay revealed a stepwise reduction of the copy number of the SRY gene from wild type control to complete SRY-KO pig compared to the GGTA1 gene. These results supported the assumption of a porcine SRY duplication.

To further exclude that these findings originated from mosaicisms, two healthy piglets were produced via SCNT using porcine fibroblasts edited with gRNAs SRY_1 and SRY_2 targeting the 5’ flanking region of the HMG domain of the SRY gene (Fig. 2b, Fig. S18) as donor cells. Sequencing of the target site revealed two deletions of 72 bp and 73 bp in both piglets (704/1 and 2) (Tab. S5, Fig. S19/S20). These results finally proved the presumptive duplication of the porcine SRY locus.

## 5 Discussion

The sex-determining region on the Y chromosome (SRY) is critically involved in mammalian male sex development (5). However, the molecular function and the role that the SRY gene plays as the main switch for male sex development in all mammals are yet to be explored. Previous studies in mice (12) and rabbits (13) investigated the potential role of the SRY gene for sex development. The murine SRY gene was knocked out by introducing two base pairs into the 5’ part of the ORF (open reading frame) of the SRY gene causing a frameshift. In one genetically male offspring this mutation led to a female phenotype (12). In rabbits, a disruption of the Sp1-binding site in the 5’ flanking region of the SRY gene also resulted in sex reversal (13). In contrast, an in-frame mutation upstream of the HMG domain of the porcine SRY gene described in this study did not result in the generation of genetically male offspring with a female phenotype. Detection of two genetic modifications in several pigs provided evidence of a presumed duplication of the porcine SRY locus. Skinner et al. described the porcine SRY gene in a two palindromic head-to-head copy manner (15), as in rabbits (16). Quantitative analysis by digital PCR (QuantStudio^®^3D, ThermoFisher Scientific) revealed duplication of the SRY locus by detection of a similar copy number of the monoallelic SRY and the biallelic GGTA1 genes. Moreover, a reduction of the copy number of the SRY gene from wild type control to a complete SRY-KO pig displayed the presumed duplication in pigs generated via intracytoplasmic microinjection. Ultimately, the generation of pigs via SCNT that carried two different deletions within the SRY gene confirmed the presence of the SRY duplication, because cloning technique avoids any mosaicism. Nanopore sequencing indicated a high similarity of the two SRY loci impairing the differentiation of both loci. The alignment of the assembled reads to the reference sequence resulted in loss of information that might be crucial for differentiation of both SRY loci. An evaluation of the raw data, a de-novo assembly or an enhancement of the Nanopore Sequencing data with Illumina MiSeq data could overcome these limitations (17-19). It is still unknown, whether both copies of the porcine SRY gene are active and required for male sex development and if there is the need to reach a certain threshold expression level from the SRY locus to induce the development of a male gender as previously described in mice (20-22). To address these questions, the identification of potential single nucleotide polymorphisms (SNPs) to differentiate between the two SRY loci is desirable.

We report here for the first time the successful knockout of the HMG domain of the porcine SRY gene by intracytoplasmic microinjection of two CRISPR/Cas9 RNP complexes resulting in genetically male pigs with a female phenotype. The CRISPR/Cas9 system has emerged as the genome editing technology of choice for many applications due to its ease of use, cost-effectivity and high specificity to introduce mutations at the targeted loci (23, 24). Nevertheless, off-target cleavages at undesired genomic sites may occur. It is necessary to further increase the specificity of the CRISPR/Cas system regarding the gRNA design (25), by involving CRISPR nickase proteins (26), using anti-CRISPR proteins (27), employing ribonucleoproteins (RNPs) (28, 29) or designing “self-restricted” CRISPR/Cas systems (30). CRISPR/Cas9 RNP components persist only temporarily in cells thereby limiting guideRNA and Cas9 expression to a short time window. The use of CRISPR/Cas9 RNPs enables efficient genome editing while significantly reducing possible off-target events and mosaicism formation. Random integration of DNA segments into the host genome as with DNA plasmids is avoided by using RNPs (28, 29, 31). However, no off-target events were found at possible sites using PCR-based analysis and Sanger sequencing in the SRY-KO pigs generated via intracytoplasmic microinjection of CRISPR/Cas RNPs. However, only with whole-genome sequencing using accurate and sensitive off-target profiling techniques such as GUIDE-Seq and CIRCLE-Seq the occurrence of unexpected mutations could be excluded completely (25, 32, 33).

In the present study, healthy SRY-KO pigs showing normal development and growth rates were born (Fig. S6, Tab. S4). Moreover, the knockout of the HMG domain resulted in piglets with a female phenotype, including female external and internal genitalia. A previous study reported that rabbits with a knockout in the Sp1-binding sites of the SRY gene showed a dramatically reduced number of follicles in their ovaries (13). Although a normal copulatory behavior was observed, no pregnancy was established by mating of the genetically modified rabbits to wild type male rabbits. Transfer of blastocysts from wild type female rabbits into pseudo-pregnant SRY-KO rabbits resulted in a successful pregnancy with the birth of twelve pups. It was assumed that the abnormal development and reduced number of follicles were responsible for the decreased fertility in the sex-reversed rabbits. In our study, substantial size differences in all female genitalia of 9-months old SRY-KO pigs compared to the age-matched wild type controls demonstrated markedly retarded development of female genitalia. It has to be clarified, whether Y chromosome induced gene and hormone expression hampered the development of female genitalia in those SRY-KO pigs. One example of the influence on female sex development from disturbed hormone profiles (androstenone and müllerian inhibition substance) in females is the bovine freemartin syndrome which leads to the masculinization of the female genitalia (34). Moreover, the absence of the second X chromosome in the SRY-KO pigs might have an impact on female sex development (35). The inactivation of one copy of the X chromosome is essential for undisturbed female development, nevertheless, several genes (mainly located on the short arm of the X chromosome) usually escape X chromosome inactivation (36). Further studies are necessary to investigate the gene expression and hormone levels in these SRY-KO pigs. An XO phenotype lacking the Y chromosome would be a promising animal model to investigate the influence of Y chromosomal gene expression and to clarify the importance of the second X chromosome in female sex development. It was previously shown, that CRISPR/Cas-mediated elimination of the murine Y chromosome is possible by targeting a cluster of genes along the Y chromosome (37). Moreover, in human embryonic stem cells it was shown that the CRISPR/Cas3 system has the potential to induce long-range chromosomal deletions (38). Both methods can be utilized to generate a porcine XO phenotype.

The results of this study further clarified the critical role of the porcine SRY gene in male sex determination. The pre-determination of sex using CRISPR/Cas9 targeting the porcine SRY gene could be of great benefit for animal welfare elimination the need for castration of male offspring to avoid boar taint by delivery of phenotypically female piglets. Currently, most piglets are surgically castrated without anesthesia shortly after birth, which raised animal welfare concerns and resulted in a ban of this practice within the EU. It was recently reported that knockout of the KISSR gene by TALEN-mediated mutagenesis resulted in the generation of boars that remained in the pre-pubertal stage lacking boar taint (39). However, to use these animals for breeding purposes, they have to be hormonally treated, which in turn might result in reduced consumer acceptance. Our results could pave the way for the production of boars that produce only female offspring by integrating a CRISPR/Cas9 vector targeting the HMG domain of the SRY gene into the porcine genome. The transgenic founder would produce feminized males (XY^SRY-^) and normal females. Alternatively, the CRISPR/Cas vector could target multiple genes on the Y chromosome during spermatogenesis to prevent development of Y-chromosomal sperm. Thereby, only female offspring would be generated. In both approches, use of a self-excising vector should result in the generation of non-transgenic offspring. It remains to be determined whether products from genome-edited animals will find market acceptance in light of a critical public debate on genome engineering in many countries. Nevertheless, the above-mentioned strategies might improve welfare in pig farming and may lead to a more sustainable pork production.

In addition to its importance for animal welfare, the SRY-KO pigs could be useful for better mechanistic insights into the human male-to-female sex reversal syndrome (Swyer syndrome) (40). Overall, 15 to 20 % of humans exhibiting male-to-female sex reversal syndrome carry mutations in or dysfunctions of the SRY gene. Most of the detected variations in humans are located within the “high mobility group” (HMG) domain of the SRY gene that is responsible for DNA binding (11, 41) and thought to act as the main functional domain for SRY protein synthesis (42-44). The murine SRY gene shows only 75% similarity to the human SRY gene. In contrast, the porcine and human SRY genes are closely related (∼85 % amino acid homology) and show similar expression profiles (5, 14). Taken this in account, the high similarity of the HMG domain and the high degree of physiological, genetic and anatomical similarity of the pig to humans renders the pig as a promising large animal model to gain insight into human sex determination and the interaction of sex chromosome related gene expression profiles (5, 45).

## 6 Materials and Methods

### Animal Welfare

Animals were maintained and handled according to the German guidelines for animal welfare and the genetically modified organisms (GMO) act. The animal experiments were approved by an external animal welfare committee (Niedersaechsisches Landesamt fuer Verbraucherschutz und Lebensmittelsicherheit, LAVES, AZ: 33.9-42502-04-17/2541), which included ethical approval of the experiments.

### Transfection of gRNAs

The CRISPR/Cas9 system was employed to induce defined deletions within the SRY gene (Ensembl transcript: ENSSSCG00000037443). Guide RNAs (gRNAs) targeting either the 5’ flanking region of the HMG domain of the SRY gene (SRY_1 and SRY_2) or encompassing the HMG box (SRY_1 and SRY_3) were designed using the web-based design tool CRISPOR (http://crispor.tefor.net/) (Fig.2). Target sequences were further analyzed via BLAST to reduce the probability for off-target events. The gRNA oligos with a BbsI overhang were cloned into the linearized CRISPR/Cas9 vector pX330 (addgene, #42230). Afterwards, two CRISPR/Cas9 plasmids were co-transfected (with a final concentration of 5 μg/μl) into male porcine fibroblasts by electroporation (Neon™ Transfection System, ThermoFisher Scientific) to test the efficacy of the plasmids to induce double-strand breaks at the targeted locus. Electroporation conditions were as follows: 1350 V, 20 mm, and two pulses. After lysis of transfected cells, the cell lysate was analyzed using SRY specific primer (SRY-F: 5’-TGAAAGCGGACGATTACAGC and SRY-R: 5’-GGCTTTCTGTTCCTGAGCAC-3’). The purified PCR product (10 ng/μl) (Invisorb^®^ Fragment CleanUp – Startec) was Sanger sequenced to detect mutations at the target site.

### In-Vitro-Fertilization and In-Vitro-Maturation

In-vitro-maturation of porcine oocytes was performed as previously described (46). Frozen boar semen from a fertile landrace boar was thawed for 30 sec. in a water bath (37 °C). The motility of sperm was analyzed using microscopy (Olympus, BH-2). After washing with Androhep^®^ Plus (Minitube) and centrifugation for 6 minutes at 600 g, approx. 75 to 100 sperm per oocyte (depending on semen capacity) were used for fertilization (no sexed sperm were utilized for fertilization). After four hours of co-incubation, the fertilized oocytes were cultured in porcine-zygote-medium (PZM-3 medium).

### Somatic cell nuclear transfer

SCNT was performed as previously described (47). Fetal fibroblasts transfected with gRNA SRY_1 and SRY_2 targeting the flanking region of the HMG domain of the SRY gene were used as donor cells. Eighty-two and eighty-six one-to two-cell embryos were surgically transferred into two hormonally synchronized German Landrace gilts (7 to 9-months old). Estrus was synchronized by application of 20 mg/day/gilt Altrenogest (Regumate^®^ 4mg/ml, MSD Germany) for 12 days, followed by an injection of 1,500 IU PMSG (pregnant mare serum gonadotropin, Pregmagon^®^, IDT Biologika) on day 13 and introduction of ovulation by intramuscular injection of 500 IU hCG (human choriongonadotropin, Ovogest^®^300, MSD Germany) 78 h after PMSG administration.

### Preparation of RNP complexes for microinjection

The Alt-R CRISPR/Cas9 system (IDT) consists of two CRISPR RNA components (crRNA and tracrRNA). The crRNA was individually designed to target the HMG domain of the SRY gene (SRY_3: 5’ – AAATACCGACCTCGTCGCAA – 3’). To generate an active gRNA, both components (crRNA and tracRNA) were annealed (95 °C for 5 min and then ramped down to 25 °C at 5 °C/min) in a ratio of 1 : 1 to reach a final concentration of 1 µg/µl. Afterwards, the gRNA complex was mixed with Alt-R S.p. Cas9 nuclease 3NLS and incubated for 10 minutes at room temperature to form an active RNP complex with a final concentration of 20 ng/µl. The second RNP complex was prepared using the individually designed synthetic single-guide RNA (SRY_1: 5’ – ATTGTCCGTCGGAAATAGTG – 3’) from Synthego. The sgRNA was mixed with purified 2NLS-Cas9 nuclease using a ratio of approximately 1 : 1.5 (0,84 µl sgRNA [25pmols] and 1.25 µl Cas9 protein [25 pmols]) and incubated for 10 minutes at room temperature. After centrifugation at 10,000 rpm for 10 minutes and 4 °C, the supernatant was transferred into a new tube. Both RNP complexes were mixed in a ratio of 1 (SRY_1) to 1.7 (SRY_3) and directly used for microinjection.

### Microinjection

The RNPs targeting the SRY gene were intracytoplasmatically co-injected into IVF-produced zygotes obtained from slaughterhouse ovaries. Therefore, approx. 10 pl of the RNP solution was injected with a pressure of 600 hPa into IVF-produced zygotes (FemtoJet, Eppendorf). The injected zygotes were cultured in PZM-3 medium at 39 °C, 5 % CO_2_ and 5 % O_2_. At day 5, when embryos reached the blastocyst stage, 31 or 32 embryos, respectively, were surgically transferred into two recipients.

### Establishing cell cultures from SRY-KO piglets

Porcine fibroblasts were isolated from ear tissue of the piglets and cultured in Dulbecco’s modified Eagle’s medium (DMEM) with 2 % penicillin/streptomycin, 1 % non-essential amino acids and sodium pyruvate and 30 % fetal calf serum (FCS) (Gibco, 10270-106). When cells reached confluency, they were lysed with EDTA/Trypsin and genomic DNA was analyzed by PCR and karyotyping.

### PCR-based genotyping

Genomic DNA of the pigs was extracted from tail tips. Cells were isolated from ear tissue. The DNA concentration was determined using the NanoDrop™ (Kikser-Biotech) system. For genotyping of the pigs, polymerase chain reaction (PCR) was employed using specific primer (SRY-F: 5’-TGAAAGCGGACGATTACAGC-3’ and SRY-R: 5’-GGCTTTCTGTTCCTGAGCAC-3’) flanking a 498 bp segment of the SRY gene (Fig. 2). PCR amplification was performed in a total volume of 50 µl : 20 ng DNA, 0.6 µM reverse and forward primer, 1.5 mM MgCl_2_, 0.2 mM dNTPs and 1.25 U Taq Polymerase. Cycling conditions were as follows: 32 cycles with denaturation at 94°C for 30 sec, annealing at 59 or 60 °C for 45 sec, extension at 72°C for 30 sec and a final extension at 72°C for 5 minutes. The standard conditions for gel electrophoresis were set up to 80 V, 400 mA and 60 min using a 1 % agarose gel. The PCR-product was purified (Invisorb^®^Fragment CleanUp-Kit, Startec) and Sanger sequenced. To further analyze the genotype of the piglets Y chromosome specific genes such as KDM6A, DDX3Y, CUL4BY, UTY, UBA1Y and TXLINGY were amplified (Supplements Table 1).

### Karyotyping of the cells

Karyotyping was accomplished on porcine fibroblasts isolated from ear tissue. After treatment of cells for 30 minutes with colcemide (Invitrogen), cells were trypsinized and metaphases were prepared according to standard procedures. Fluorescence R-banding using chromomycin A3 and methyl green was performed as previously described in detail (48). At least 15 metaphases were analyzed per offspring. Standard karyotype of the pig includes 38 chromosomes. Karyotypes were described according to Gustavsson, 1988 (49) and the International System for Human Cytogenetic Nomenclature (ISCN).

### Histology

Porcine ovarian tissues were fixed with 4 % paraformaldehyde for 6 to 8 hours (smaller tissues of up to 5 × 10 mm) or overnight (tissues of up to 2 × 3 cm), incubated in 30 % sucrose for two hours and frozen at - 80 °C. Afterwards, the tissues were embed in TissueTek^®^ (Sakura, TTEK), cut in thin sections (15 μm) and stained with hematoxylin and eosin (HE) following standard procedures (50). Analyzes of inner structure of ovaries were done by microscopy (DMIL LED, Leica).

### Off-target analysis

The top ten off-target effects were selected from the gRNA design tool CRISPOR (http://crispor.tefor.net/). PCR primers used for amplifying the PCR product are listed in Supplements Table 3 for SRY_1 and Supplements Table 4 for SRY_3. The PCR product was purified (Invisorb^®^Fragment CleanUp-Kit, Startec, Germany) and analyzed via Sanger sequencing.

### DigitalPCR

Three assays including a probe and two primers (in a ratio of 2.5 probe to 9 nM primer) targeting the SRY and KDM6A genes (FAM™ -labeled) on the Y chromosome and GGTA gene (HEX™ -labeled) on chromosome 1 (as control) were designed (IDT) for digital polymerase chain reaction (dPCR). The dPCR was performed in a total reaction volume of 14.5μl with the following components: 7.3 μl Master Mix (QuantStudio™ 3D Digital PCR Master Mix v2, ThermoFisher Scientific), 0.7 μl HEX™ and VIC™ dye-labeled assays each, 1.4 μl diluted genomic DNA and 4.4 μl nuclease-free water. Standard dPCR thermal cycling conditions were used with an annealing temperature of 60 °C in the QuantStudio™ 3D Digital device (ThermoFisher Scientific). Copy numbers of the genes within each chip were compared and analyzed via the QuantStudio™ 3D AnalysisSuite software (http://apps.lifetechnologies.com/quantstudio3d/). The copy number of the GGTA1 gene was set at 2 (biallelic), copy numbers of KDM6A and SRY genes were given in proportion to the GGTA1 gene. All findings were verified in three replicates with variable DNA concentration and different samples (51).

### Nanopore Sequencing

Whole genome sequencing was performed by using the MinION device of Oxford Nanopore Technologies to investigate the porcine SRY locus. DNA from a male wild type blood sample (2 ml) was purified with the NucleoBand^®^HMW DNA Kit (Macherey-Nagel). To eliminate fragments below 40 kb the Short Read Elimination Kit XL (Circulomics) was utilized. Subsequently, 47 μl high molecular weight DNA (30 - 40 ng/μl) was prepared with the Ligation Sequencing Kit (SQK-LSK109, Oxford Nanopore) and the NEBNext^®^ Companion Module for Oxford Nanopore Technologies^®^ Ligation Sequencing (BioLabs, E7180S) using the Nanopore Oxford standard protocol for ligation sequencing.

## 7 Acknowledgements

This research did not receive any specific grant from funding agencies in the public, commercial, or not-for-profit sectors. Gudrun Göhring received a research grant from RIBIRTH. All other authors disclose any financial and personal relationships with other people or organizations that could inappropriately influence this work.

The authors are grateful to the IVF and SCNT team, Petra Hassel, Maren Ziegler, Roswitha Becker and Antje Frenzel for their effort in producing the SRY-KO pigs. We thank the staff from the pig facility for taking care of the pigs. We also thank Dr. Lutz Wiehlman for the cooperation and support to perform Nanopore Sequencing in the Research Core Unit Genomics in the MHH and Collin Davenport for the assembly of the data.

The pX330-U6-Chimeric_BB-CBh-hSpCas9 was a gift from Feng Zhang (Addgene plasmid # 42230; http://n2t.net/addgene:42230; RRID:Addgene_42230).

