## Supplemental Data for "The knockout of the HMG domain of the porcine SRY gene causes sex reversal in gene-edited pigs"

**10 Supplementary information**

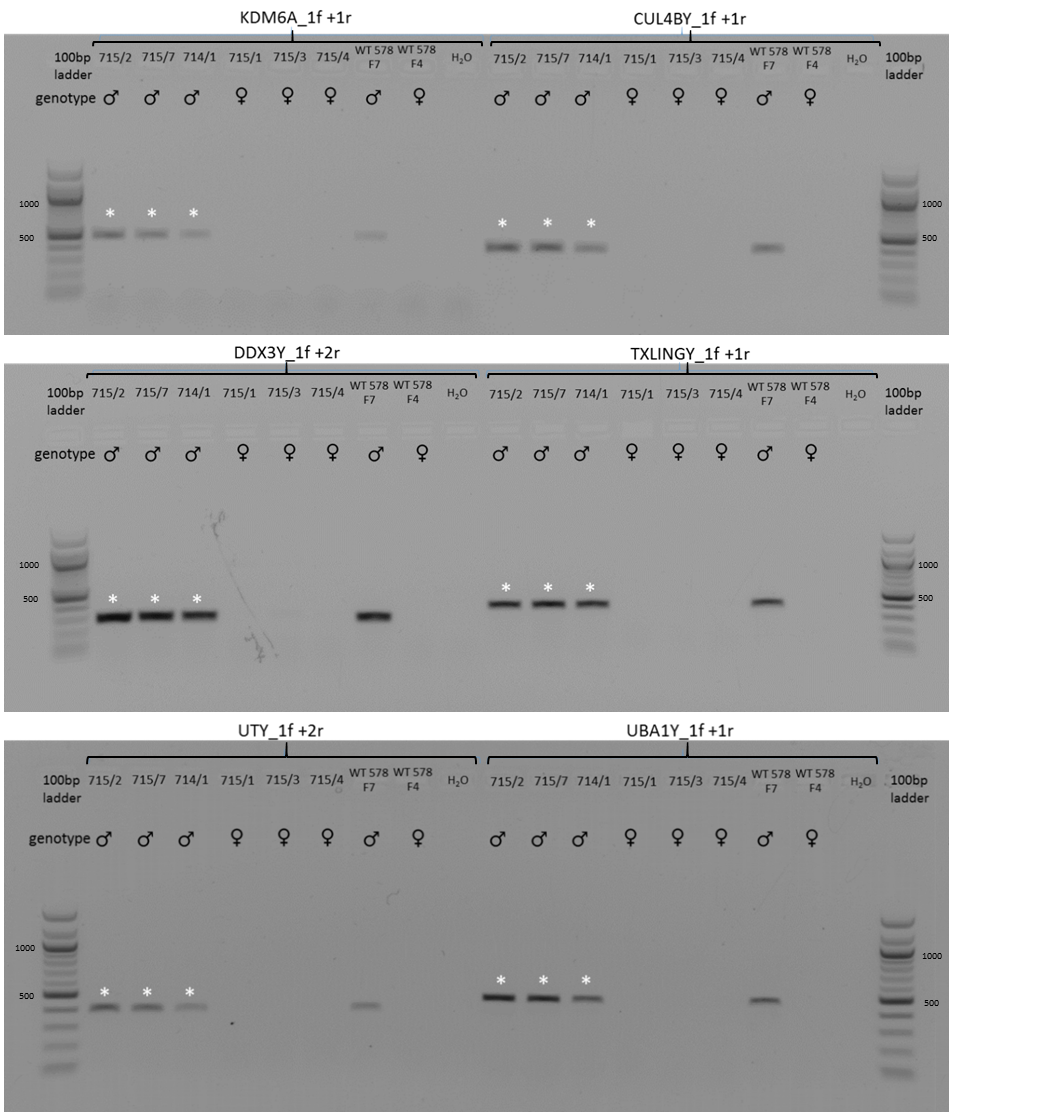

**Figure S1** PCR of six different Y chromosome specific genes (KDM6A, CUL4BY, DDX3Y, TXLINGY, UTY and UBA1Y) for detection of the Y chromosome in SRY-KO piglets (715/2, 715/7 and 714/1, indicated by a white asterisk) compared to female wild type controls (715/1, 715/3 and 715/4) from same litter. Moreover, a male (WT 578/F7) and a female (WT 578/F4) DNA sample was used as positive and negative control.

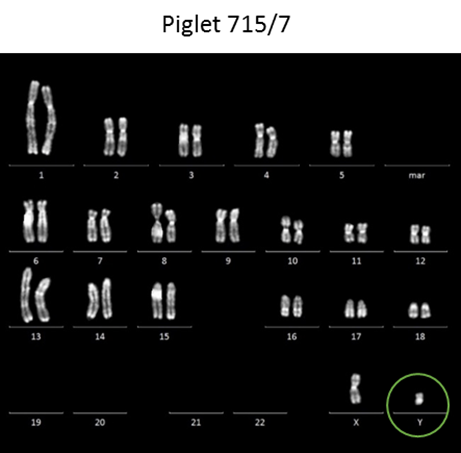

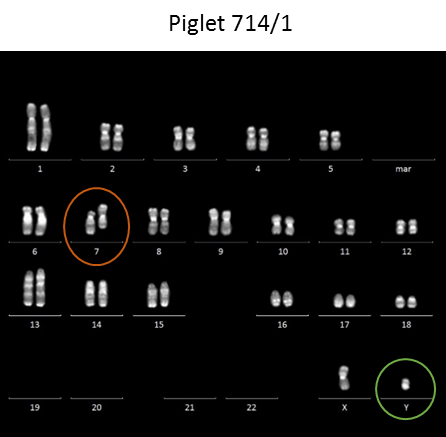

**Figure S2** Karyotyping of the SRY-KO piglet 715/7 and 714/1 confirming the male genotype by analysis of the sex chromosomes. In piglet 714/1, a clonal aberration (inversion) could be found in chromosome 7.

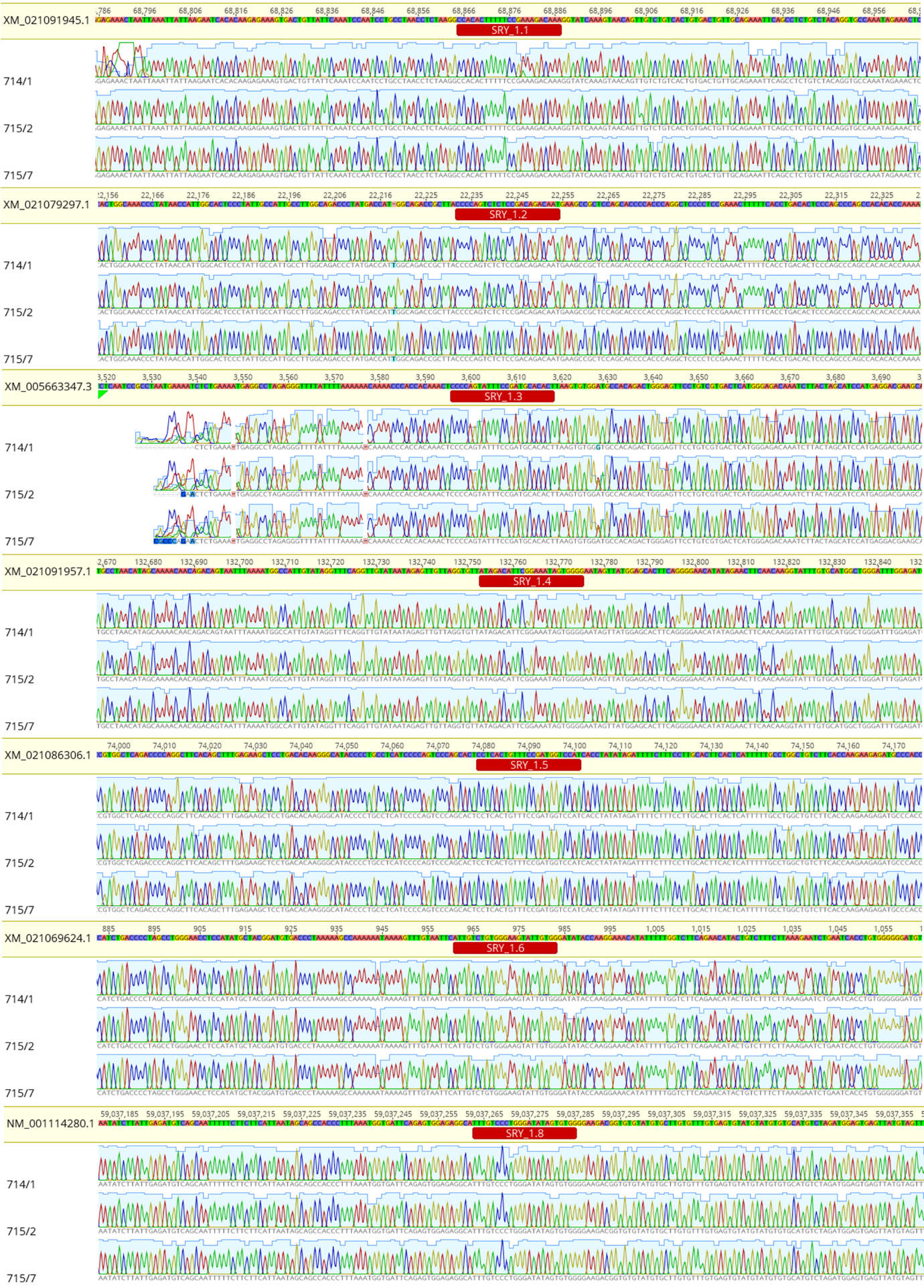

**Figure S3** Sanger Sequencing for detecting potential off-target sites of gRNA SRY_1 in SRY-KO pigs 714/1, 715/2 and 715/7. Overall, no off-target events were found.

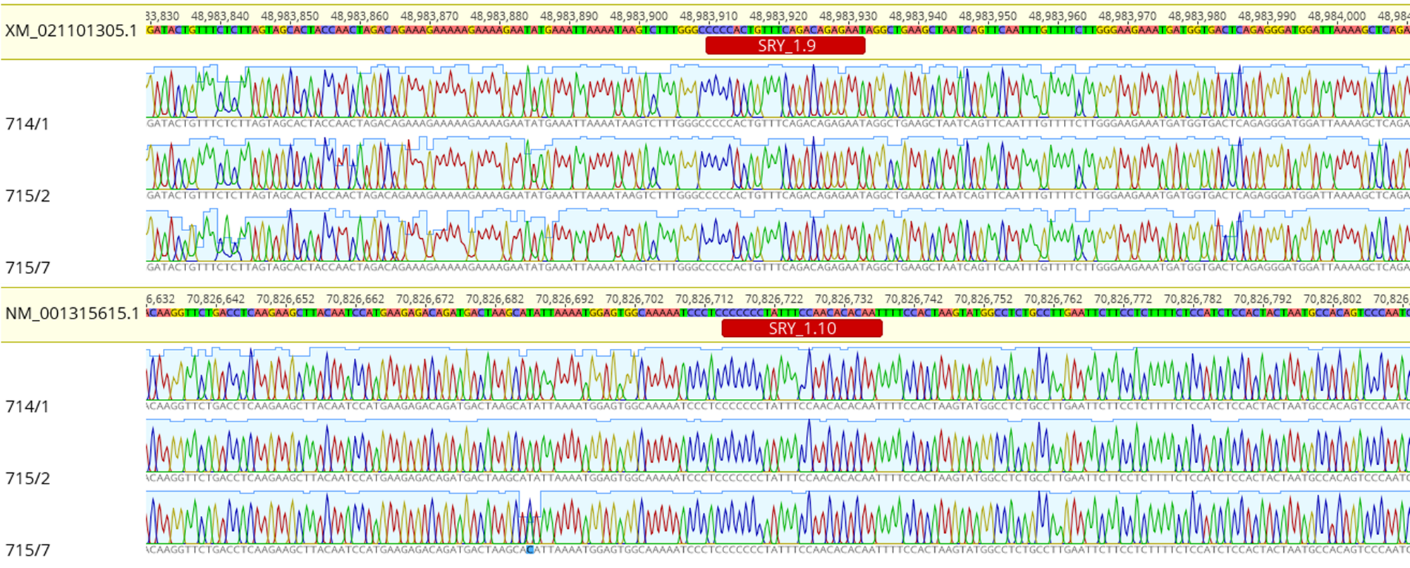

**Figure S4** Sanger Sequencing for detecting potential off-target sites of gRNA SRY_1 in SRY-KO pigs 714/1, 715/2 and 715/7. Overall, no off-target events were found.

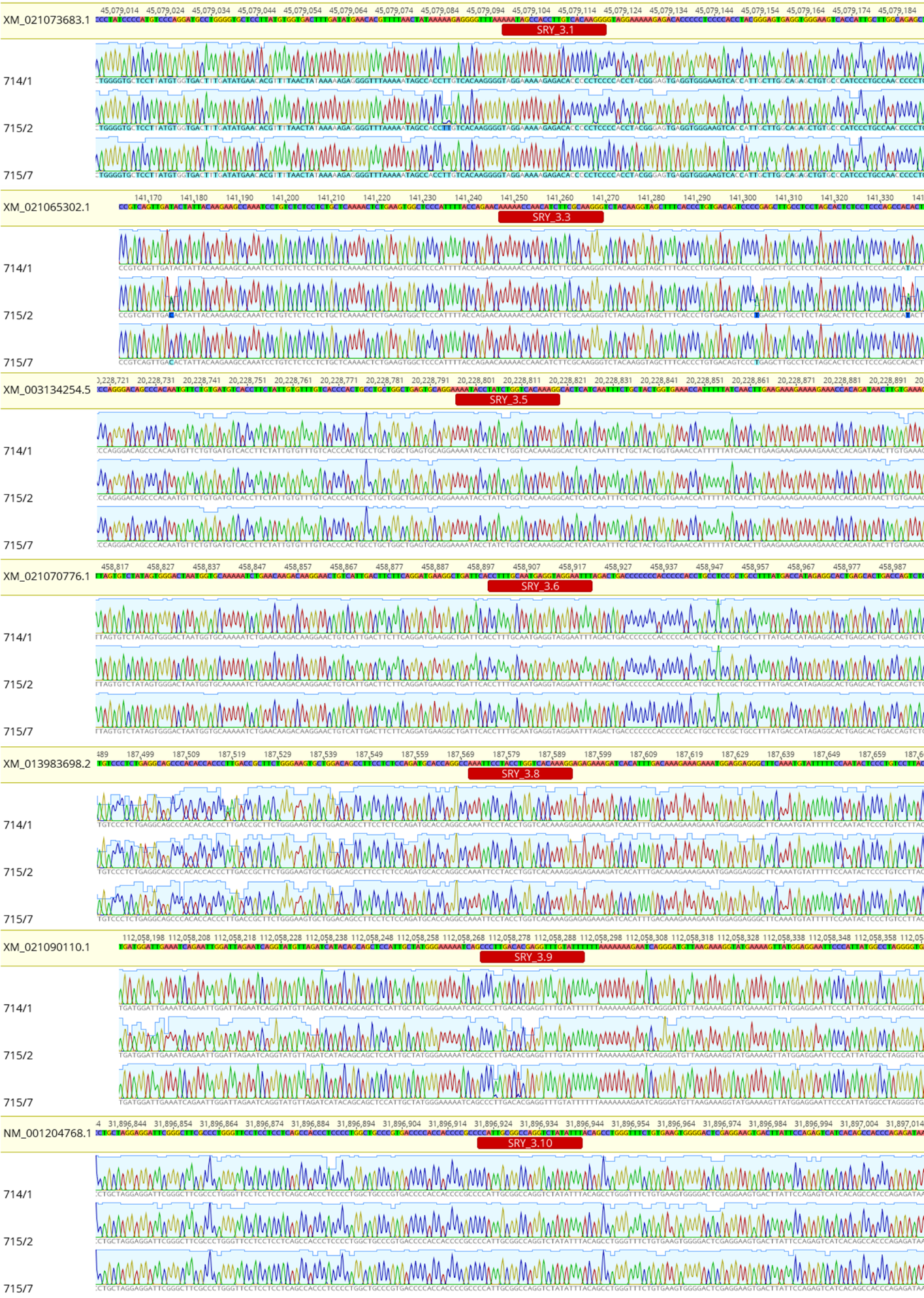

**Figure S5** Sanger Sequencing for detecting potential off-target sites of gRNA SRY_3 in SRY-KO pigs 714/1, 715/2 and 715/7. No off-target events were found.

Growth rate (WT vs. SRY-KO pigs)

Weight (kg)

| Age (weeks) | WT male | WT female | MI WT female | SRY-KO (715/7) | SRY-KO (714/1) |
| --- | --- | --- | --- | --- | --- |
| 1 | 2.85 | 3 |  |  |  |
| 2 | 4.75 | 5.8 | 4.9 | 4.4 | 5.2 |
| 3 | 7.5 | 8.3 | 6.1 | 6.9 | 7 |
| 4 | 7 | 8.2 | 10 | 9.7 | 10.9 |
| 5 | 8.25 | 8.5 | 10 | 10 | 11 |
| 6 | 10.75 | 11 | 10.5 | 13 | 15 |
| 7 | 13 | 13.7 | 15 | 17.5 | 20.5 |
| 8 | 15.5 | 15.25 | 19.1 | 21.9 | 24.5 |

**Figure S6** Schematic diagram displaying the weight development (median of growth rates) of the SRY-KO pigs 715/7 (yellow) and 714/1 (dark blue) compared to the male (light blue) and female WT controls (orange) and female littermates (MI WT females, grey) with the age of 1 to 8 weeks. The weights of pigs were determined once per week and the median of growth rates per group was calculated to illustrate the diagram. WT males (n)= 6; WT females (n)= 5; MI WT females (n)= 7

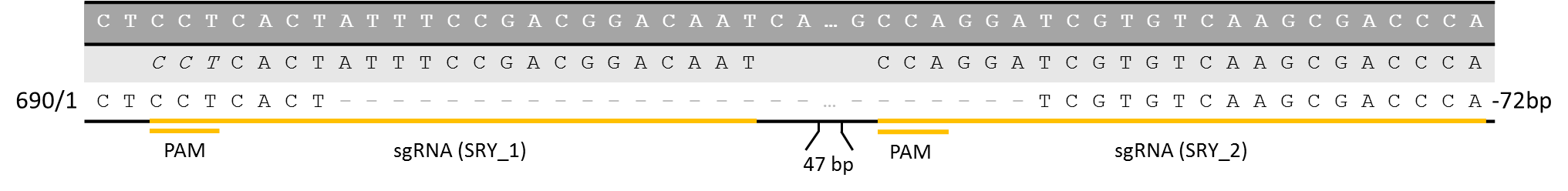

**Figure S7** Sanger sequencing of piglet 690/1 revealed a 72 bp deletion within the 5’ flanking region of the HMG domain of the SRY gene.

**A**

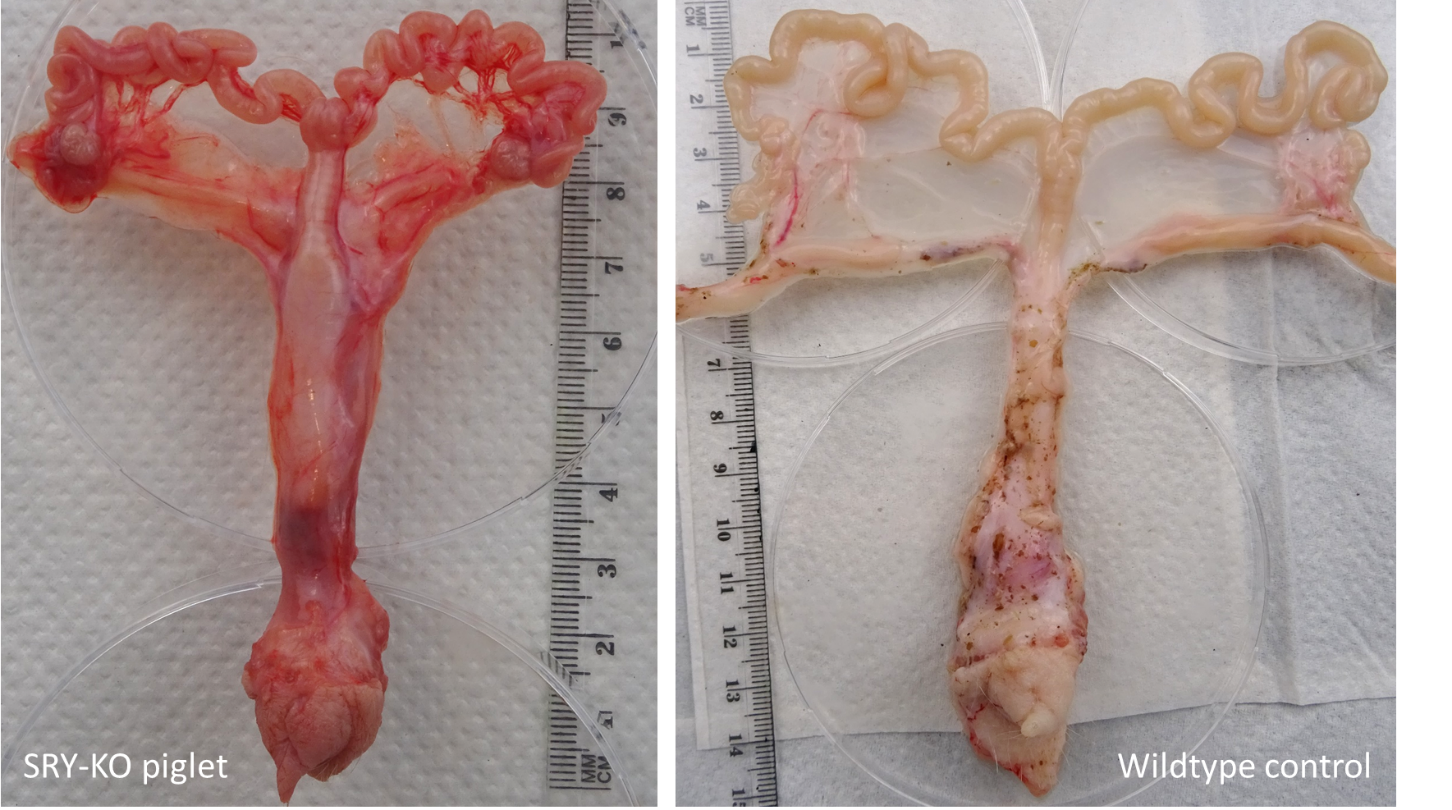

**B**

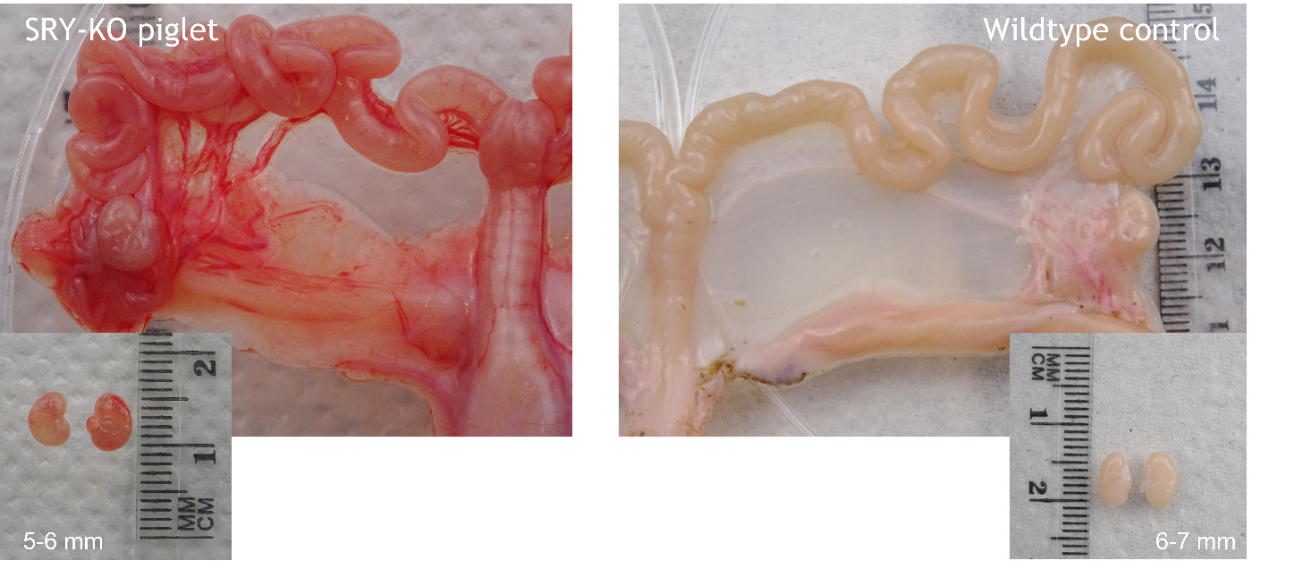

**Figure S8** The uteri, oviducts and ovaries of the SRY-KO, XY piglet (715/2) and the WT,XX piglet (control from artificial insemination) at day 34. **a** No differences were shown in size of oviduct and uteri. **b** Solely the ovaries in SRY-KO, XY piglets are approx. 2-fold smaller than the ovaries of the WT, XX piglet.

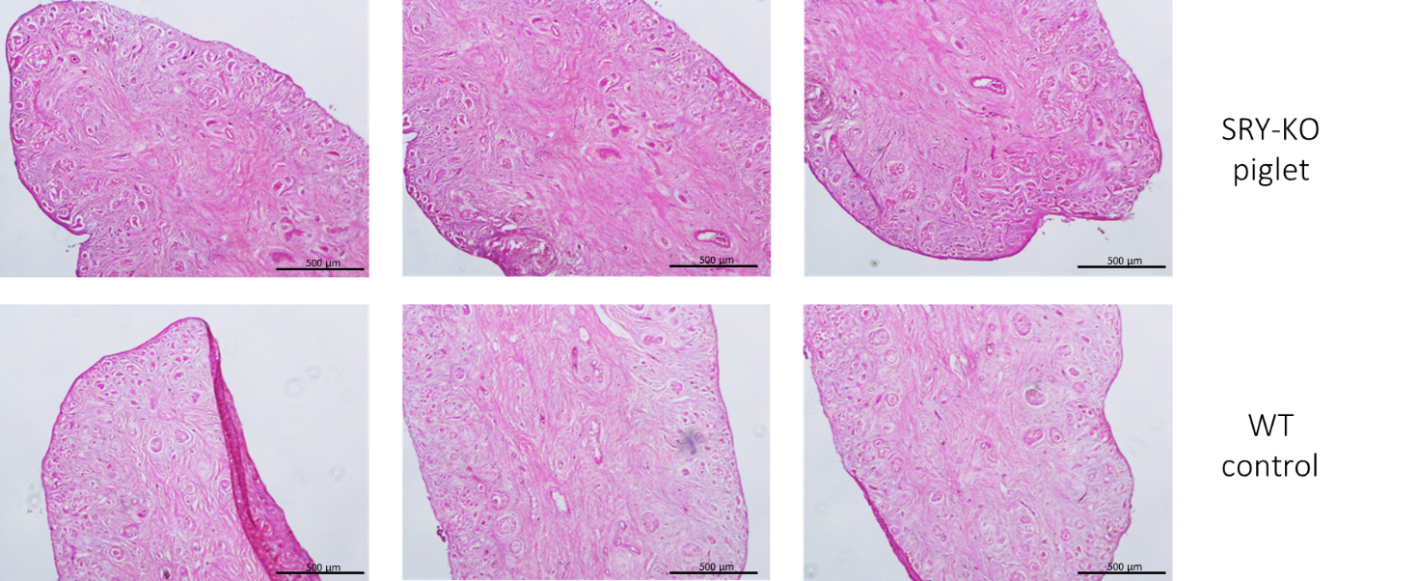

**Figure S9** H&E-Staining of porcine ovarian tissue from the SRY-KO piglet (upper images) und wild type (WT) control (lower images) 34 days after birth. No structural differences were shown.

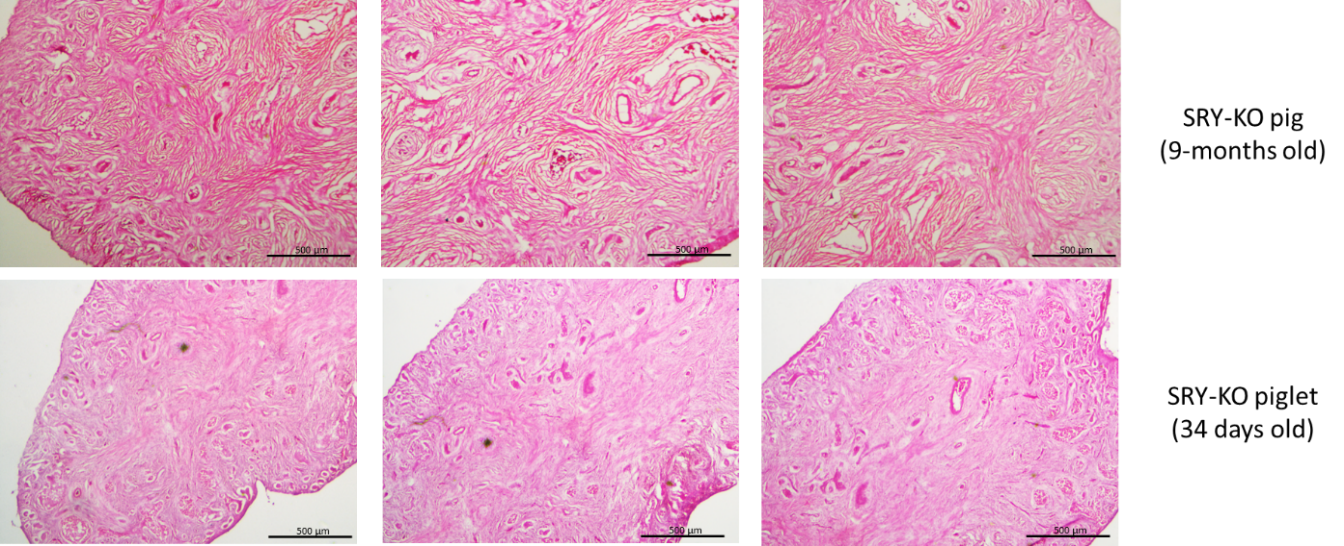

**Figure S10** Histological analysis of the ovarian tissue of the 9-months old SRY-KO pig (upper images) compared the SRY-KO piglet at the age of 34 days (lower images). A higher amount of loose connective tissue in the 9-months old SRY-KO pig indicated fat deposits within the ovarian tissue compared to the 34 days old SRY-KO piglet.

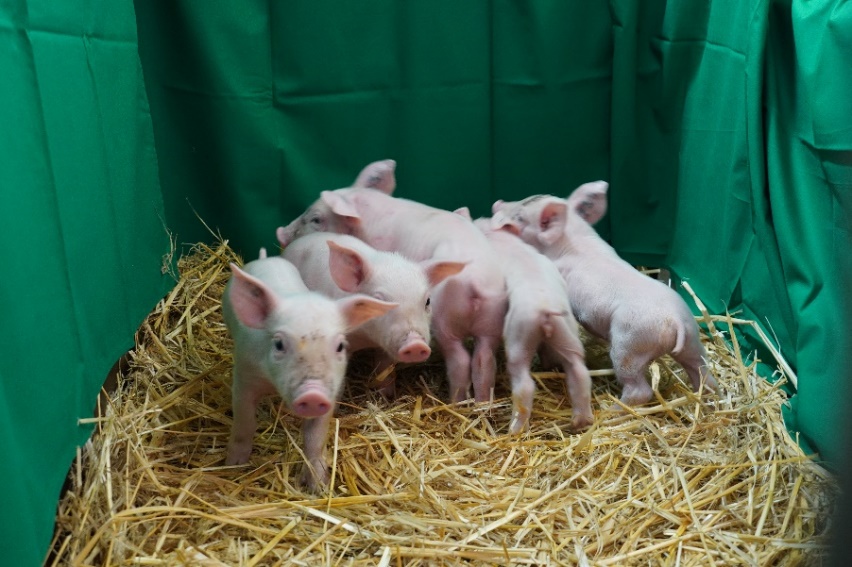

**Figure S11** Healthy piglets with a female phenotype and male genotype were born from re-cloning of cells from the SRY-KO piglet 715/2.

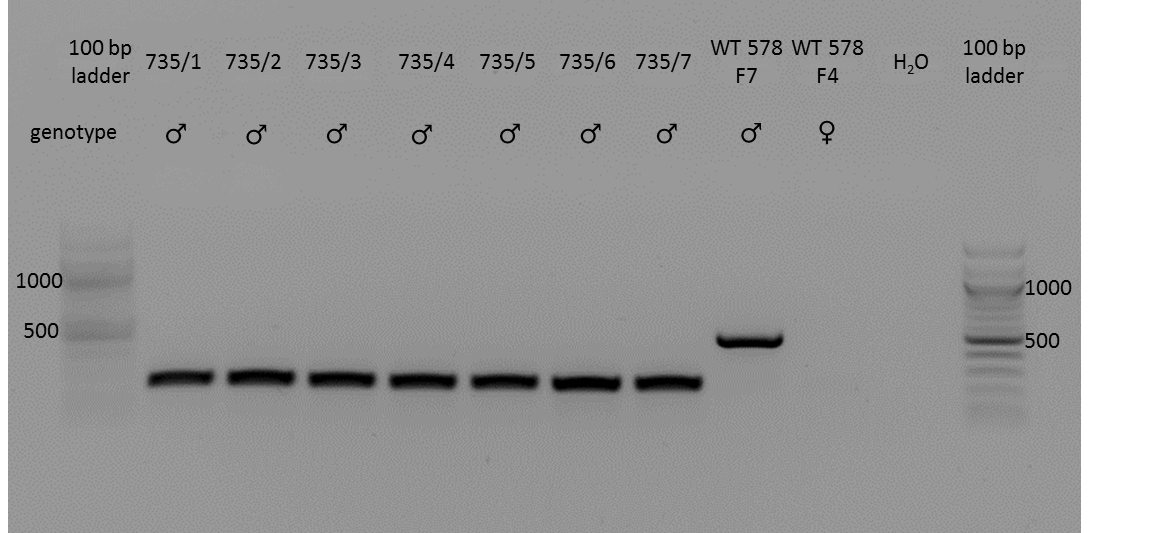

**Figure S12** PCR of piglets from re-cloning of cells from the SRY-KO piglet 715/2. All piglets showed deletions of ~300 bp within the SRY gene (indicated with white asterisks) compared to the expected band of approx. 500bp in male wild type control (WT 578 F7).

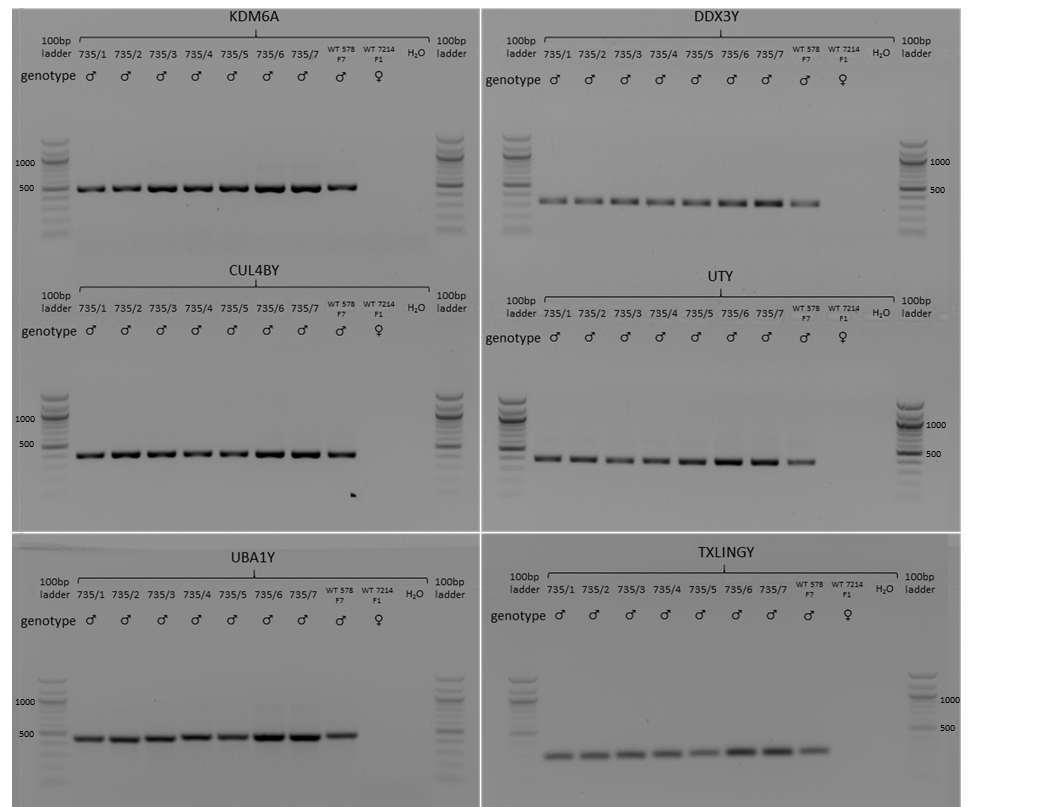

**Figure S13** Detection of six different Y chromosome specific genes (KDM6A, CUL4BY, DDX3Y, TXLINGY, UTY and UBA1Y) via PCR in SRY-KO piglets from re-cloning (735/1-7). Male (WT 578/F7) and female (WT 7214/F1) DNA samples was used as positive and negative control.

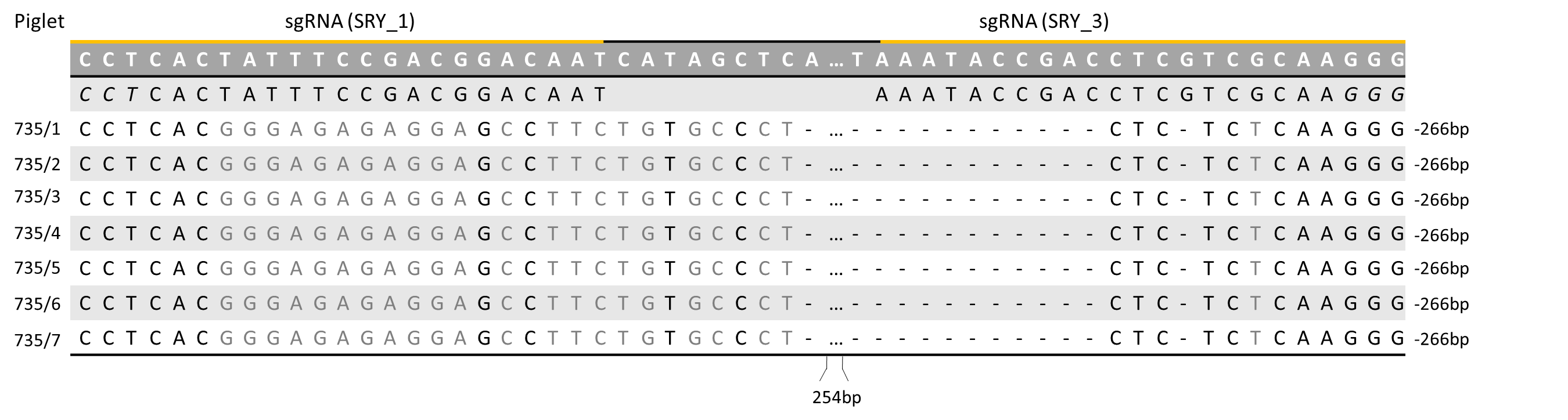

**Figure S14** Sanger sequencing revealed a deletion of -266bp in all of the piglets generated via re-cloning of the SRY-KO piglet 715/2.

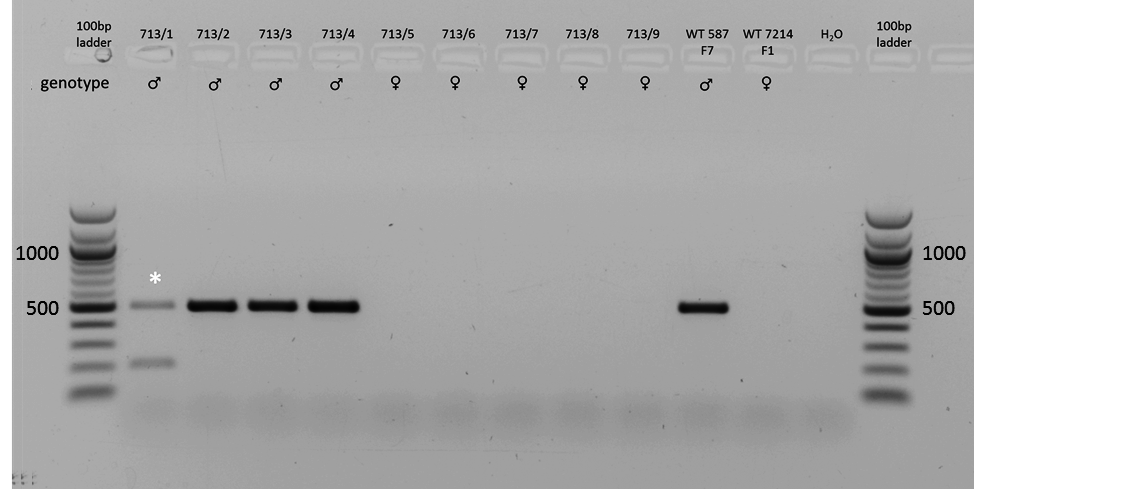

**Figure S15** PCR-based detection of the SRY gene in piglet 713/1-9. Five pigs showed a female geno- and phenotype, while four displayed a male geno- and phenotype. No sex reversal occurred. One pig (713/1) revealed two different genetic modification within the SRY gene (indicated with a white asterisk). A male (WT 587/F7) and a female (WT 7212 F1) DNA sample served as positive and negative control.

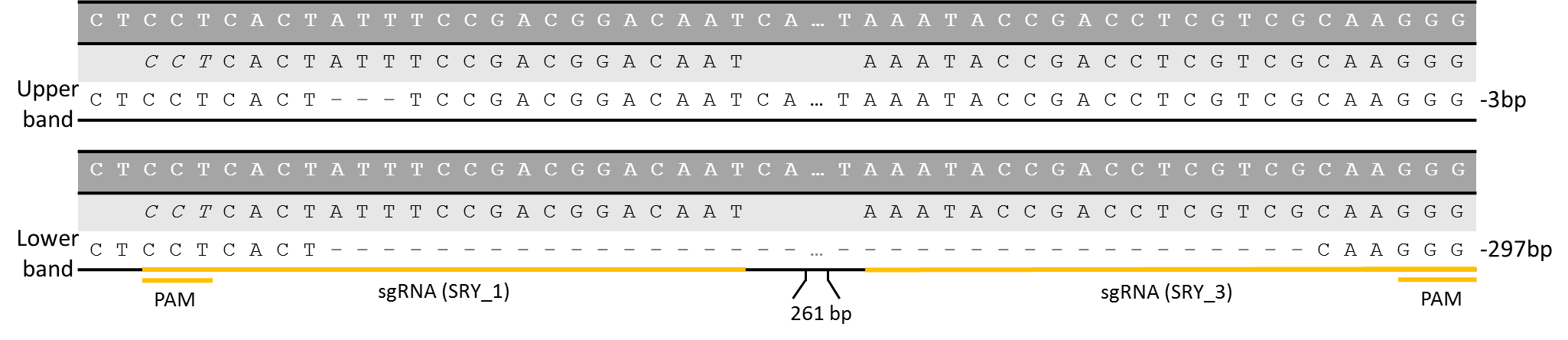

**Figure S16** Sanger sequencing of DNA isolated from cells of piglet 713/1 revealed two genetic modification, including a 3 bp and 297 bp deletion within the SRY gene.

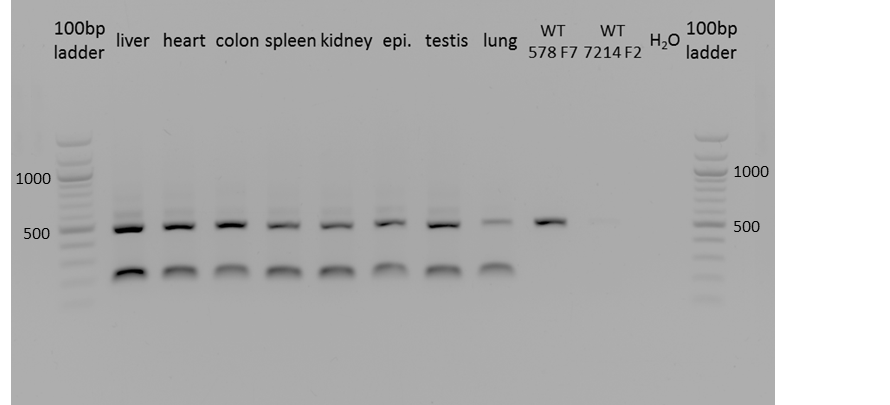

**Figure S17** PCR-based detection of genetic modifications within the SRY gene in different organ samples (liver, heart, colon, spleen, kidney, epididymis, testis and lung) from piglet 713/1. All DNA samples revealed two genetic modifications (further confirmed by Sanger sequencing). A male (WT 578 F7) wild type control served as positive control and WT 7214 F2 (female) as negative control.

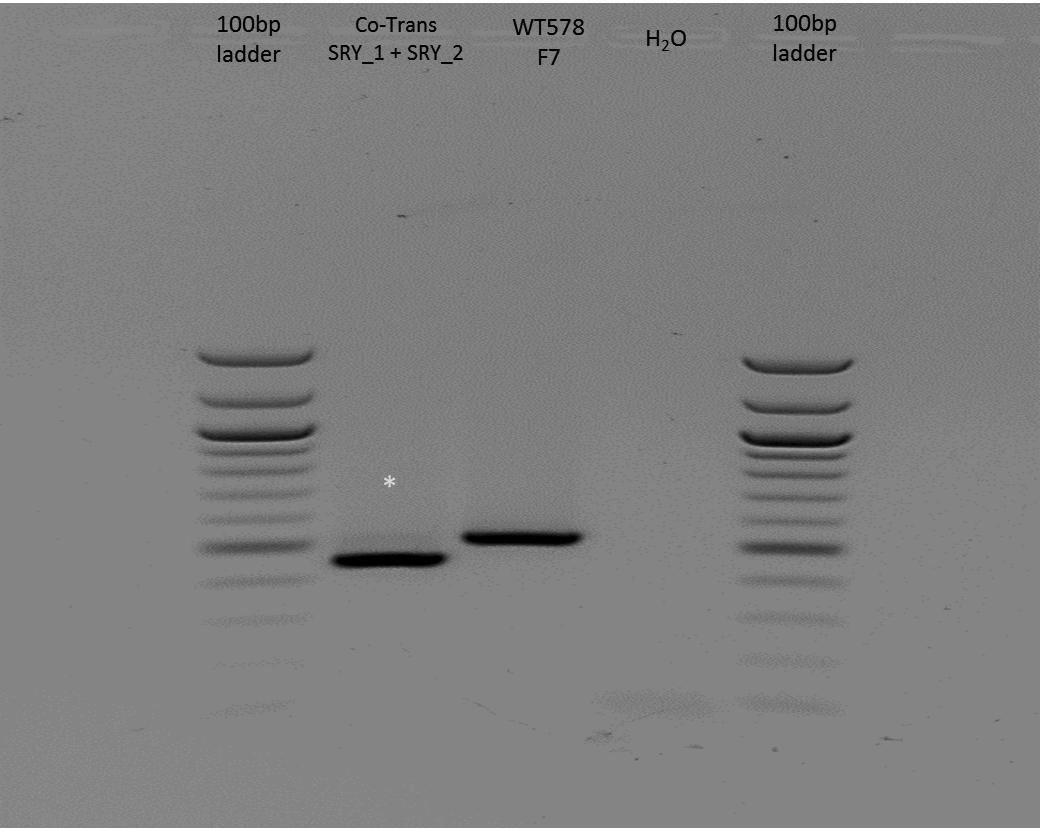

**Figure S18** PCR after co-transfection of two plasmids (SRY_1 and SRY_2) in male fetal fibroblasts and selection of the edited cells via single cell dilution (lower band). The edited cells revealed a mutation of approx. 70bp (indicated with a white asterisk) compared to WT control. The male WT control (WT 578 F7) showed an expected band of approx. 500bp. The edited cells were further employed for somatic cell nuclear transfer.

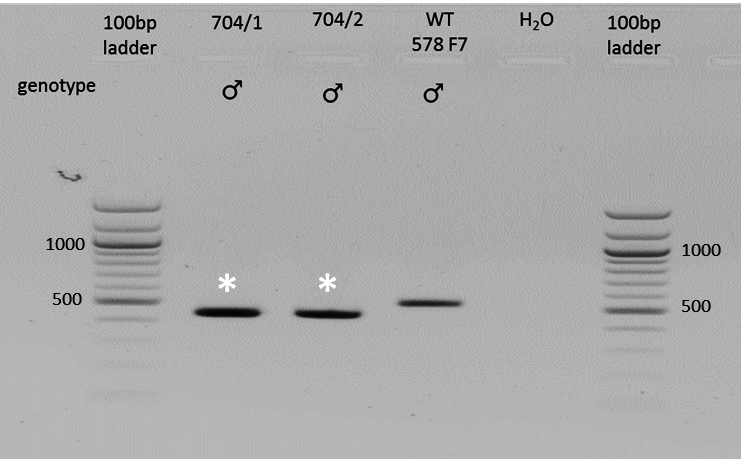

**Figure S19** PCR to detect genetic modifications in the SRY gene of the two piglet (704/1-2) generated via SCNT. Both piglets showed a mutation of approx. 70bp compared to the male WT control (WT 578 F7). The WT control displayed an expected band of ~500bp.

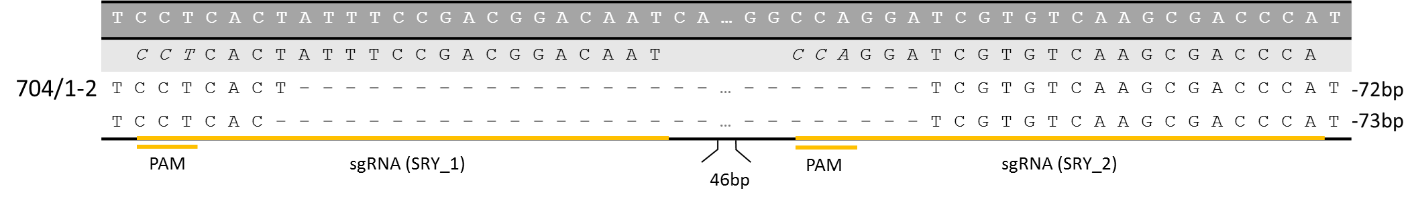

**Figure S20** Both piglets (704/1 and 704/2) generated via SCNT of two gRNAs (SRY_1 and SRY_2) targeting the 5’ flanking region of the HMG box of the SRY gene showed two different mutation including a deletion of -72bp and -73bp.

**Table S1** For Y chromosome detection, six different Y chromosome specific genes (KDM6A, CUL4BY, DDX3Y, TXLINGY, UTY and UBA1Y) were utilized.

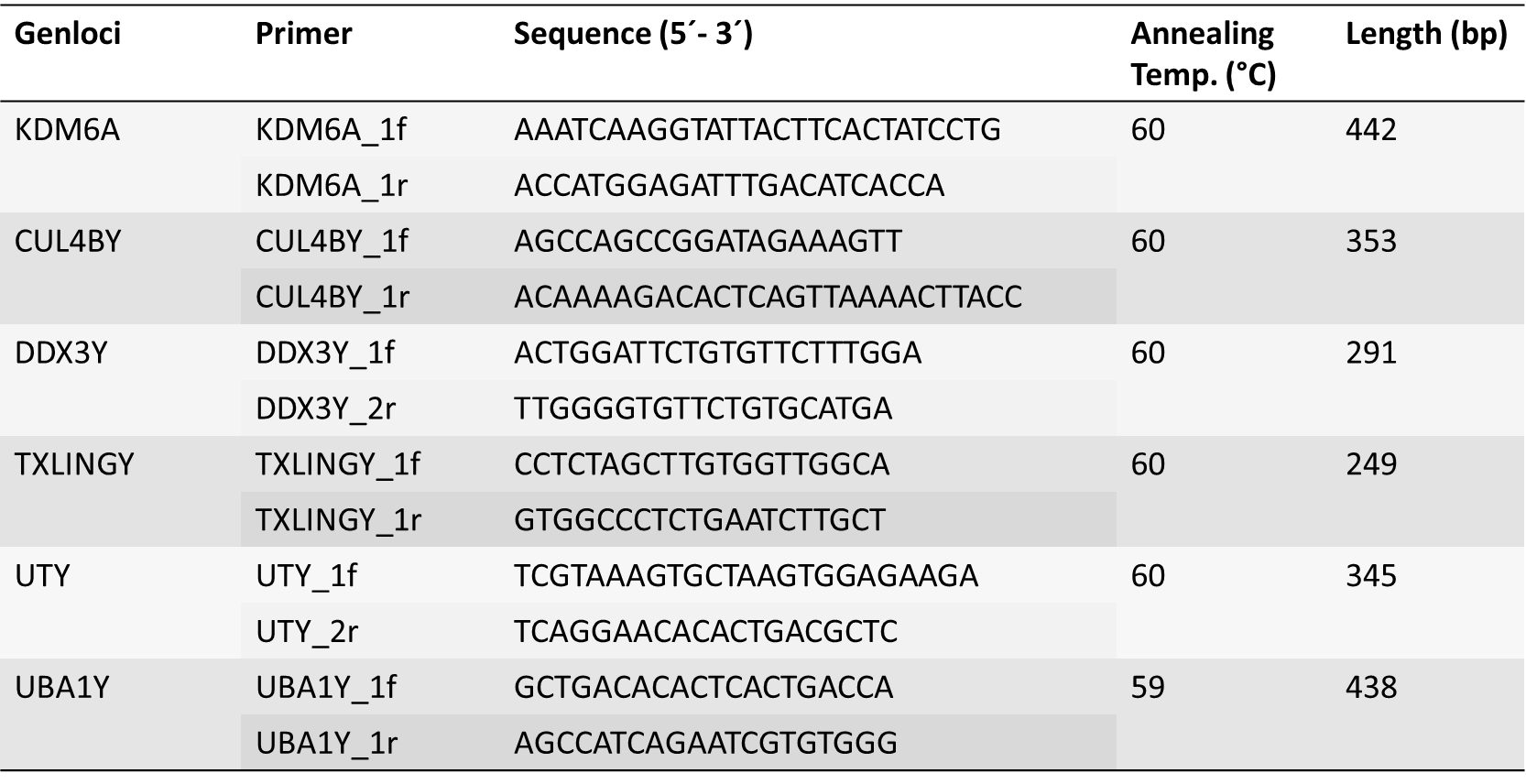

**Table S2** Top ten off-target sites for gRNA SRY_1 on the porcine genome. Primer pair for sequencing of the off-target events are listed. One off-target site could not be amplified.

| **Genloci** | **Primer** | **Sequence (5’ – 3’)** | **Annealing**  **Temp. (°C)** | **Length (bp)** |
| --- | --- | --- | --- | --- |
| Chr. 5 - intergenic:  XM_021091945.1 | SRY_1.1_f | GAGGCTGAACTGGGAACCTT | 60 | 2,509 |
|  | SRY_1.1_r | AGCACATGCTCTCTGCCAAA |  |  |
| Chr. 18 - intergenic:  XM_021079297.1 | SRY_1.2_f | CATGCGCAGTCTGAACAAGG | 58 | 3,134 |
|  | SRY_1.2_r | GGATGAAGGGTGCTAGACGG |  |  |
| Chr. 4 - intron:  XM_005663347.3 | SRY_1.3_f | GAGCGCATCAACTGAGTGAC | 65 | 1,227 |
|  | SRY_1.3_r | GGAAATGAGACAGGCCACCT |  |  |
| Chr. 5 - intron:  XM_021091957.1 | SRY_1.4_f | CTCTGACTTGCACCCTGCTT | 62 | 2,131 |
|  | SRY_1.4_r | ACTTCTCAATCCGCCCTATGC |  |  |
| Chr. 3 - exon:  XM_021086306.1 | SRY_1.5_f | AACAACATGCGTCCAAACCG | 62 | 1,776 |
|  | SRY_1.5_r | GCATCAGCACTCACCTGGAT |  |  |
| Chr. 13 - exon:  XM_021069624.1 | SRY_1.6_f | AGTGACTGGGTTTGGGGTTG | 62 | 2,002 |
|  | SRY_1.6_r | CGCCAGAGTCCCATACACTC |  |  |
| Chr. 4 - intergenic:  XM_021090127.1 | SRY_1.7_f | - | - | - |
|  | SRY_1.7_r | - |  |  |
| Chr. 1 - intergenic:  NM_001114280.1 | SRY_1.8_f | CCTGACCACGTCGTATCTCG | 61 | 3,035 |
|  | SRY_1.8_r | AGCTAAGGGTGGAGTTTGGC |  |  |
| Chr. 8 - intergenic:  XM_021101305.1 | SRY_1.9_f | TAGCACCCCCAGAAACTCCT | 62 | 2,602 |
|  | SRY_1.9_r | GGTGGATACTGTCAGCTGGG |  |  |
| Chr. 16 - intergenic:  NM_001315615.1 | SRY_1.10_f | CCTAATTTGGCCTGCGCTTC | 61 | 3,134 |
|  | SRY_1.10_r | ACCTCTGAGGGTGTGACCTT |  |  |

**Table S3** Top ten off-target events for gRNA SRY_3 on the porcine genome. Primer pair for sequencing the off-target sites are listed. Three off-target sites could not be amplified.

| **Genloci** | **Primer** | **Sequence (5’ – 3’)** | **Annealing Temp. (°C)** | **Length (bp)** |
| --- | --- | --- | --- | --- |
| Chr. 14 - intergenic:  XM_021073683.1 | SRY_3.1_f | CTCCCCACAGCTGCTCTTTT | 62 | 2,487 |
|  | SRY_3.1_r | GAATTGGGCACTTGCTGGAC |  |  |
| Chr. 12 - intergenic:  XM_003358165.4 | SRY_3.2_f | - | - | - |
|  | SRY_3.2_r | - |  |  |
| Chr. 11 - intergenic:  XM_021065302.1 | SRY_3.3_f | AACAGGGAACCATCCACCAA | 61 | 3,463 |
|  | SRY_3.3_r | CTCCAGGAGGCCATATGCTG |  |  |
| Chr. 1 - intergenic:  XM_021093786.1 | SRY_3.4_f | - | - | - |
|  | SRY_3.4_r | - |  |  |
| Chr. 17 - intergenic:  XM_003134254.5 | SRY_3.5_f | AGCCTTATCCAATGAGGCCG | 66 | 2,866 |
|  | SRY_3.5_r | CTAATGCCAGGGCAGTTTGC |  |  |
| Chr. 13 - intron:  XM_021070776.1 | SRY_3.6_f | CAAAAGGCTACCAGGGGTGT | 61 | 2,418 |
|  | SRY_3.6_r | GCCCAAGGTGACCTCAAACT |  |  |
| Chr. 13 - intron:  XM_021069725.1 | SRY_3.7_f | - | - | - |
|  | SRY_3.7_r | - |  |  |
| Chr. 14 - intergenic:  XM_013983698.2 | SRY_3.8_f | CCCTGCCGTTAGATCCAGTC | 66 | 4,663 |
|  | SRY_3.8_r | TCCAGAGGGCACCTGTGATA |  |  |
| Chr. 4 - intron:  XM_021090110.1 | SRY_3.9_f | CTTCTCTGGTAACTGGCCCC | 66 | 2,382 |
|  | SRY_3.9_r | TCCCGCAGATCCATTCCAAC |  |  |
| Chr. 3 - intergenic:  NM_001204768.1 | SRY_3.10_f | CTTGTTCCTTCCTGGGTGGG | 62 | 2,269 |
|  | SRY_3.10_r | CTCCAGATGGGGGACACTTG |  |  |

**Table S4** Overview of the weight development (mean, median and standard derivation of growth rates) of the SRY-KO pigs (715/5 and 714/1) compared to the male and female WT controls and female littermates (MI WT females) with the age of 1 to 8 weeks. The weights of pigs were determined once per week and the mean, median and standard derivation of growth rates per group calculated. WT males (n)= 6; WT females (n)= 5; MI WT females (n)= 7;

|  | **WT males** | | | **WT females** | | | **WT MI females** | | | **SRY-KO**  **(715/7)** | **SRY-KO**  **(714/1)** |
| --- | --- | --- | --- | --- | --- | --- | --- | --- | --- | --- | --- |
| **Weight (kg) per week** | Mean | Median | Standard deviation | Mean | Median | Standard deviation | Mean | Median | Standard deviation |  |  |
| 1. Weeks | 2.85 | 2.85 | 0.19 | 2.94 | 3 | 0.65 | 4.87 | 4.9 | 0.58 | - | - |
| 2. Weeks | 4.68 | 4.75 | 0.86 | 5.26 | 5.8 | 1.14 | 6.74 | 6.1 | 1.04 | 4.4 | 5.2 |
| 3. Weeks | 7.22 | 7.5 | 1.45 | 7.76 | 8.3 | 1.42 | 10.2 | 10 | 0.87 | 6.9 | 7 |
| 4. Weeks | 7.02 | 7 | 1.56 | 7.74 | 8.2 | 1.36 | 10.31 | 10 | 0.9 | 9.7 | 10.9 |
| 5. Weeks | 7.75 | 8.25 | 1.65 | 8.2 | 8.5 | 1.5 | 11.79 | 10.5 | 2.74 | 10 | 11 |
| 6. Weeks | 10.17 | 10.75 | 1.79 | 10.1 | 11 | 1.36 | 15.93 | 15 | 3.26 | 13 | 15 |
| 7. Weeks | 12 | 13 | 2.3 | 12.76 | 13.7 | 1.87 | 19.51 | 19.1 | 3.82 | 17.5 | 20.5 |
| 8. Weeks | 14.42 | 15.5 | 2.32 | 14.75 | 15.25 | 2.02 | 21.23 | 20 | 4.66 | 21.9 | 24.5 |

**Table S5** Transfer of embryos generated by SCNT of cells transfected with CRISPR/Cas plasmids SRY_1 and SRY_2 into two recipients. Two piglets were born. Both of them showed a male phenotype and no sex reversal.

| Recipient | Transferred embryos | Pregnancy | Offspring | Genetic modification on the SRY gene | Sex reversal |
| --- | --- | --- | --- | --- | --- |
| 7263 (704) | 82 | + | 2 | 2 | - |
| 7266 | 86 | - | - | - | - |
